# Dopamine reward transients calibrate movement timing via one-shot updates to behavioral vigor

**DOI:** 10.1101/2025.04.04.647288

**Authors:** Allison E Hamilos, Isabella C Wijsman, Qinxin Ding, Pichamon Assawaphadungsit, Zeynep Ozcan, Elias Norri, Kimberly Reinhold, Bernardo L. Sabatini, John A Assad

## Abstract

Dopamine neurons (DANs) play seemingly distinct roles in reinforcement,^1–3^ motivation,^4,5^ and movement,^6,7^ and DA-modulating therapies relieve symptoms across a puzzling spectrum of neurologic and psychiatric symptoms.^8^ Yet, the relationship among these roles is unknown. Here, we show DA’s tripartite functions are causally linked by a process in which phasic striatal DA rapidly and persistently recalibrates the propensity to move, a measure of vigor. Using a self-timed movement task, we found that single exposures to reward-related DA transients (both endogenous and exogenously-induced) exerted one-shot updates to movement timing—but in a surprising fashion. Rather than reinforce specific movement times, DA transients quantitatively *changed* movement timing on the next trial, with larger transients leading to earlier movements (and smaller to later), consistent with a stochastic search process that calibrates the frequency of movement. Both abrupt and gradual changes in external and internal contingencies—such as timing criterion, reward content, and satiety state—caused changes to the amplitude of DA transients that causally altered movement timing. The rapidity and bidirectionality of the one-shot effects are difficult to reconcile with gradual synaptic plasticity, and instead point to more flexible cellular mechanisms, such as DA-dependent modulation of neuronal excitability. Our findings shed light on how natural reinforcement, as well as DA-related disorders such as Parkinson’s disease, could affect behavioral vigor.

## MAIN TEXT

Both electrical^9^ and optogenetic activation^10–12^ of dopamine neurons (DANs) can gradually reinforce actions.^10–15^ It is widely believed DANs drive gradual learning via phasic dopamine (DA) transients, with ongoing debate over what these transients represent in reinforcement learning frameworks (e.g., causal association,^16^ learning rate,^3^ value,^4,5^ reward/performance-prediction errors^1,2,17–22^, etc.). Yet, multiple lines of evidence suggest DA also plays “real-time” roles in motor control^6,7,23–26^ and motivation,^5,25^ which are thought to be distinct from DA’s function in gradual learning.^6,22,24,27^ For example, tonic DA levels are associated with movement vigor, with both tonic DA and vigor increasing in reward-rich environments.^4–6,27^ Exogenous DA/DAN manipulations also causally modulate the probability,^7,23,24^ number,^23^ acceleration^23,24^ and variability^11^ of movements, and DA-modulating drugs are used ubiquitously in clinical settings as a “tuning dial” to adjust patients’ propensity to move.^8,28^ DA repletion/DA-receptor agonists increase movement propensity (e.g., to treat Parkinson’s disease), whereas DA-receptor antagonists have the opposite effect (e.g., to treat mania, Huntington’s disease, delirium, etc.). Yet, it remains a mystery whether *endogenous*, reward-related DA transients are related to DA’s function in modulating the propensity to move.^23,27,29–31^

Because DA drugs bidirectionally influence the propensity to move, we suspected DA’s roles in reinforcement and movement might be mechanistically linked by a tuning process, whereby endogenous DA transients calibrate the propensity to move based on feedback from recent reward history. Individual DANs encode both reward-and movement-related information,^31,32^ suggesting these functions may be intertwined or somehow co-dependent.

We recently showed that DANs play a causal, real-time role in movement initiation in mice performing a self-timed movement task.^7^ Intriguingly, in similar timing paradigms, hypokinetic conditions (e.g., Parkinsonism/DA-receptor antagonists) cause systematically late movement timing compared to control subjects,^33–36^ whereas hyperkinetic conditions (e.g., DA-receptor agonists) cause early timing.^37,38^ These findings indicate that movement timing can provide an intuitive, quantitative definition of “propensity to move” (earlier timing=higher propensity; later timing=lower). Furthermore, because subjects receive correct/incorrect feedback after each trial of the self-timed movement task, they can update their timing on the next trial to improve their chances of receiving reward, allowing us to relate outcome-related DA signaling to how subjects calibrate their propensity to move based on the timing requirements of the task.

### Task for quantifying propensity to move

We trained head-fixed, water-controlled mice to perform a self-timed movement task^7^ (**Fig 1a**). On each trial, mice had to estimate elapsed time to decide when to lick after a brief cue (tone/LED). Mice received a juice reward if the first lick occurred during a hidden reward window (3.3-7s after the cue), which had to be discovered through trial-and-error. Only the first lick determined the trial outcome (unrewarded/rewarded). Correctly timed first-licks triggered an immediate correct tone, whereas those that occurred too early triggered an error tone, and failure to lick before the end of the reward window triggered a distinct miss tone.

**Fig 1.**
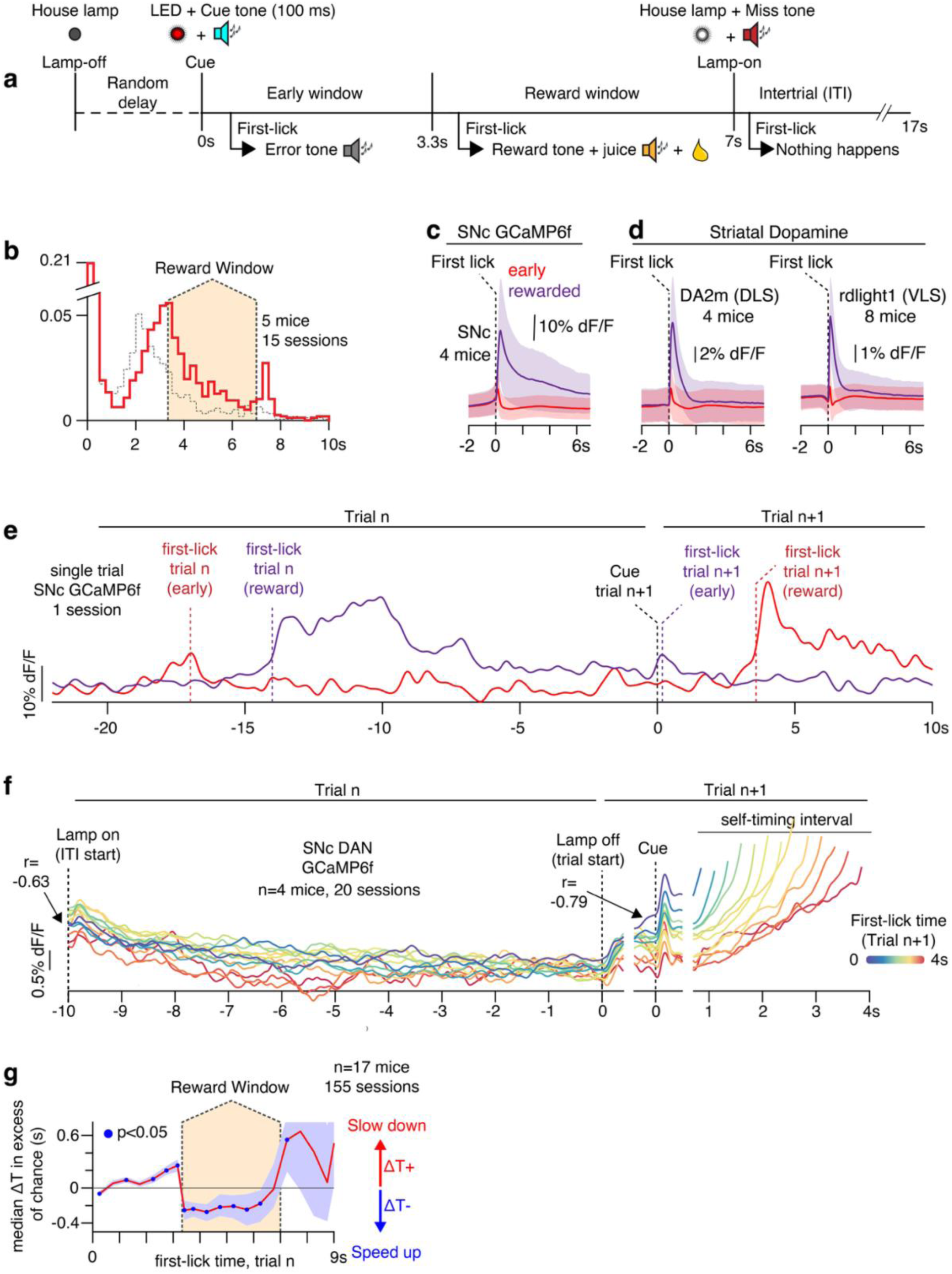
Outcome-related DA is related to timing on the next trial. a, Self-timed movement task. **b,** Example timing distributions, pooled among same 5 mice as in Fig 4. Black distribution: beginning of session (trials 1-125); Red distribution: trials 376-500. **c-d,** Average outcome-related DAN/DA signals; shading: std. **e-f,** Timing-related DAN signals: **e,** Single examples of an early (red) and rewarded (purple) trial on trial n. **f,** DAN signals pooled by movement time and plotted up to 150ms before spout contact. X-axis breaks indicate change of alignment. **g,** Median change in timing on trial n+1 as a function of first-lick time on trial n (all mice, all sessions), corrected for potential regression to the median effects (see Methods). Rewarded trials were followed by systematically earlier timing, whereas unrewarded trials (early and late) were followed by later timing. Shading: 95% CI (10,000x bootstrap).

Mice learned to self-time licks, producing timing distributions with broad variability typical of natural timing behavior (**Fig 1b**).^7,33,39–42^ The peak of the timing distribution reproducibly occurred at or before the earliest possible rewarded time, even in well-trained animals, consistent with findings from previous primate and rodent timing studies^7,40–43^ (**Extended Data Fig 1a-b)**. In our task, we found this bias was due to even well-trained animals initially favoring early movement times at the beginning of each session (**Fig 1b** dashed trace) before gradually shifting movement times toward the reward window later in the session (**Fig 1b** red trace). These reproducible behavioral patterns presented a unique opportunity to observe the relationship between DAN signaling and timing updates. We first exploited the natural variability of the timing distributions to ask whether DAN signaling was related to trial-to-trial updates in movement timing. We then examined whether DAN signaling could explain longer-term trends in timing behavior, which could serve as a potential calibration mechanism for the propensity to move.

### Outcome-related DA signals

We used fiber photometry to monitor intracellular Ca^++^ in genetically-defined DANs expressing GCaMP6f, recording from both cell bodies (substantia nigra, SNc: n=4 DAT-cre mice; ventral tegmental area, VTA: n=3 mice) and axon terminals (dorsolateral striatum, DLS, n=4 mice). DANs exhibited RPE-like transients in the timing task: unrewarded first-licks were followed by small dips in intracellular Ca^++^; rewarded trials were followed by large increases (**Fig 1c**). These signals were surprisingly persistent, decaying for >10s after the reward, much longer than GCaMP6f’s off-kinetics (t½ <300 ms at 37°C),^44^ and lasting seconds past cessation of the lick-bout, indicating they were not due to licking movements *per se*. The same basic RPE-like features were present across DAN recording sites (SNc, VTA, DLS) and in striatal DA release (**Fig 1d**), which we monitored with fluorescent DA sensors (dLight or GRAB-DA2m), expressed in either dorsolateral (DLS) or ventrolateral dorsal striatum (VLS^45,46^; **Extended Data Fig 2**).

By themselves, the persistent RPE-like signals in somatic DAN Ca^++^ are perhaps unlikely to exert downstream effects. For example, axonal DAN Ca^++^ and striatal DA signals were much less persistent than the somatic DAN Ca^++^ signals in **Fig 1d**. Nonetheless, they point to a potential inter-trial interaction: The persistent somatic DAN Ca^++^ signals bled into the next trial (**Fig 1e**), becoming continuous with differences in baseline DAN Ca^++^ signals. These baseline signals were in turn correlated with timing on the *next* trial (**Fig 1f**; baseline signal averaged over –0.4 to 0s before cue onset; Pearson’s r: –0.79, 95% CI: [-0.84,-0.73]). Moreover, these systematic baseline offsets could be traced back to—and were correlated with—the average amplitude of the DA transient at the end of the *previous* trial, such that larger DA transients were associated with higher subsequent baseline signals and relatively early movement times on the next trial, whereas smaller DA transients (and dips) were associated with lower subsequent baseline signals and relatively late movements (**Fig 1f)**. Because the amplitude of the DA transient on the previous trial depended largely on whether that trial had been rewarded or not (**Fig 1c**), these combined observations suggest that 1) the trial outcome might influence future timing; and 2) the amplitude of the DA transient might mediate this putative trial-to-trial timing update. We address these hypotheses in turn:

### Relationship of previous trial outcome to movement time on next trial

There were four possible trial outcomes: 1) early (no reward), 2) rewarded, 3) late (no reward, during the ITI), or 4) no lick. Because mice only rarely licked too late or did not lick, we initially focused analyses on early versus rewarded outcomes. Intuitively, an early (unrewarded) trial should prompt slower timing on the next trial to increase the chances of being rewarded next time. Conversely, following a rewarded trial, we might expect animals to move at about the same time on the next trial to receive reward again. To examine this, we compared the change in the movement time between consecutive trials, ΔT_(n+1)-n_. A potential confound is that, on average, relatively late trials might—by chance—tend to be followed by earlier trials (and relatively early followed by later). This “regression to the median” phenomenon could occur even in the absence of goal-directed updates to timing. We thus compared the change in the movement time between consecutive trials, ΔT_(n+1)-n_, to the change in movement time expected by chance from regression to the median (n=17 mice, 155 sessions; see **Methods**). We found that ΔT_(n+1)–n._ was consistently in excess of chance (**Fig 1g**). After unrewarded trials, animals indeed moved significantly later on the next trial (update after early trials: +74ms, 95% CI: [+41ms, +106ms]; 10,000x bootstrap procedure). However, surprisingly, after rewarded trials, rather than move at the same time on the next trial, animals moved significantly *earlier* (update after rewards: –317ms, 95% CI: [–406ms, –223ms]). That is, after a rewarded trial, rather than repeating that rewarded movement-time, mice *sped up* on the next trial.

These combined effects suggest a “sloshing”-like phenomenon, whereby timing updates tended to adjust back and forth between earlier and later lick-times, depending on whether the animal was rewarded or unrewarded on the previous trial. This may seem unintuitive (and inconsistent in some respects with normative reinforcement-learning (RL) algorithms; see Discussion), but it is not necessarily irrational: A thirsty mouse may want to move at the earliest possible rewarded time, but this time is uncertain—mice had to discover it through experience. By slowing down after early trials and speeding up after rewarded trials, this sloshing effect can cause the timing distribution to build up around the earliest possible rewarded time.

### DA transients explain *changes* in timing

We next examined whether DA signals could explain these timing updates. We derived a generalized linear model (GLM) predicting the *change* in the movement time on the next trial, ΔT_(n+1)-n_, from the amplitude of outcome-related DA transients (**Fig 2a**). This amplitude was encoded as the DA signal averaged over 0-500ms following the previous trial’s first-lick (“DA”, **Fig 2b**). We controlled for optical artifacts and ongoing body movements by including the tdTomato (“tdt”) photometry channel, accelerometer and/or neck EMG signals as predictors.

**Fig 2.**
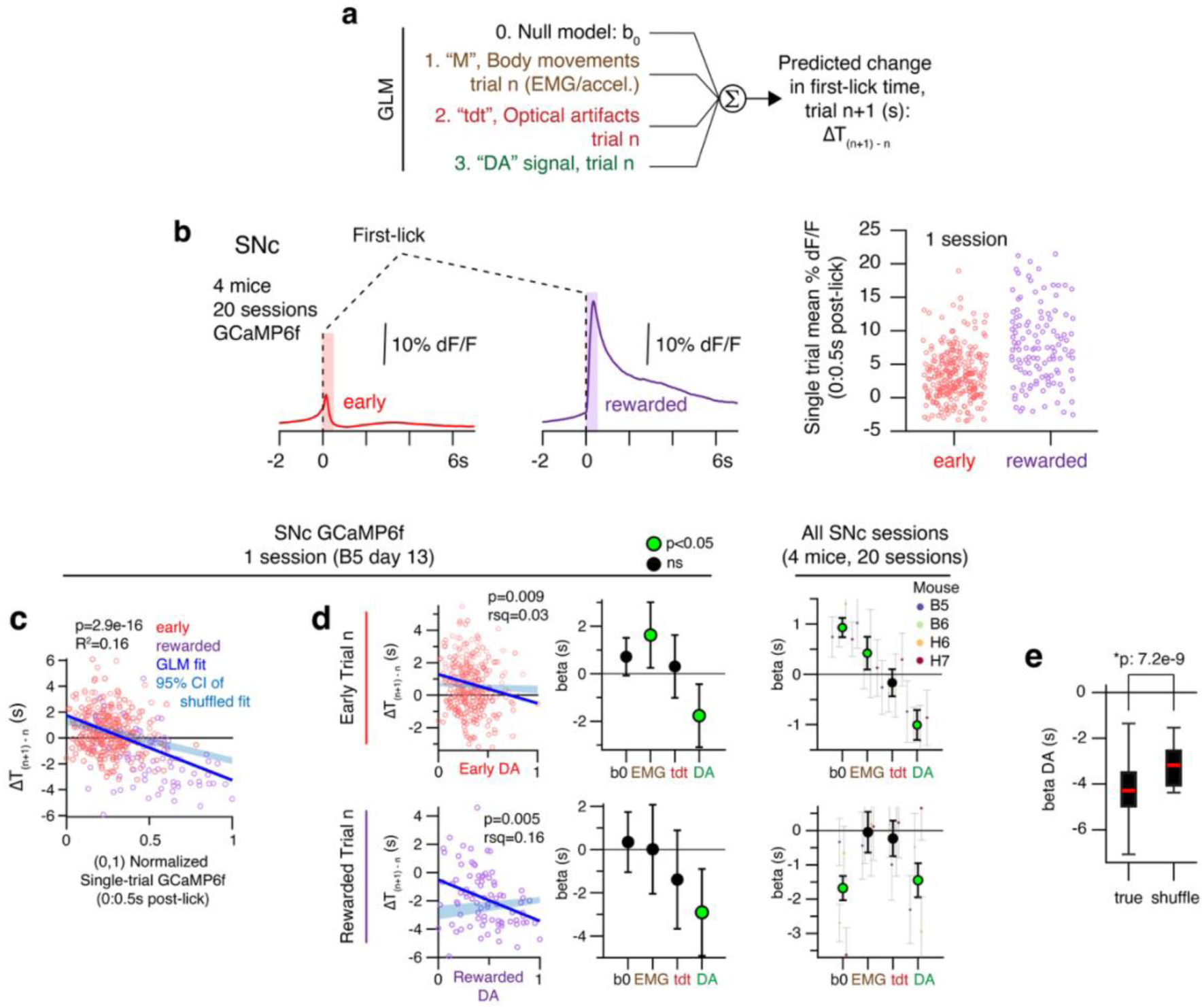
Outcome-related DAN signals predict changes in timing on the next trial. a, GLM. **b,** Left: average outcome-related DAN signals. Right: trial-by-trial variability in single-trial DAN signals. Each point is from a single trial, with signal averaged over a 0.5s window after the first-lick (shaded rectangles in Left panel). Points are jittered horizontally for clarity. **c-d,** GLM fit. Panel *c* shows model fit for 1 session. Panel *d* shows the same model fit for each trial outcome separately. Blue shading in scatter plots shows the range of fit coefficients obtained by chance from shuffled models (10,000 fits on data with shuffled trial order). Coefficient error bars: 95% CI. **e,** Distribution of model DA coefficients from all 20 sessions, true data vs 10,000x shuffle fit. Standard box plot. *p: Wilcoxon rank sum test. Model fits for each DAN/DA recording modality and site shown in **Extended Data Figs 3-4**.

Across all recording sites (and for both GCaMP and DA sensors), DA was independently predictive of ΔT_(n+1)–n_ (**Fig 2c**; **Extended Data Fig 3**), in excess of optical artifacts and ongoing body movements. The larger the DA transient amplitude, the larger the timing update on the next trial (highest DA: largest early-shifts; lowest DA: largest late-shifts).

The amplitude of the DA transient was strongly correlated with trial outcome (which was also a strong predictor of ΔT_(n+1)–n_). Although this would not preclude a causal role for DA, we tested if additional information was present in DA by fitting separate models for early and rewarded trials. Both early and rewarded DA transient amplitudes showed considerable trial-by-trial variability (**Fig 2b**), which had additional predictive power beyond correlation with trial outcome (**Fig 2d, Extended Data Fig 3**). These effects were also not explained away by a tendency to revert toward the median movement time (DA coefficient significantly larger than expected by chance in shuffled trial order model (p=7.2e-9, Wilcoxon rank sum test; **Fig 2e**; **Extended Data Fig 4**).

Outcome-related DA was thus positioned to make “one-shot” learning updates to timing. We use the machine learning term “one-shot learning” here because a *single exposure* to an endogenous DA transient was associated with a quantitative update to behavior. To our knowledge, DA learning effects based on variability in endogenous DA reward transients have not previously been observed.

### DAN stimulation causes one-shot timing updates

To test if outcome-related DA could cause one-shot timing updates, we virally expressed channelrhodopsin (ChR2, n=4 DAT-cre mice) in genetically defined DANs for optogenetic manipulations (**Fig 3a**). To modulate outcome-related DAN signaling, we triggered stimulation in the SNc at the time of the first-lick on 30% of randomly interleaved trials. Activation with ChR2 was calibrated^7,31^ both outside the context of the task and within the task to recapitulate endogenous DA reward transients (**Fig 3b**), which matched the amplitude and width of unstimulated reward transients during the task (**Fig 3c; Extended Data Fig 5a**). Activation on rewarded trials slightly increased DA transient amplitude, but on average, this remained within the limits of unstimulated reward transients on single trials measured during the same session (**Extended Data Fig 5c**). High power blue-light stimulation causes photoswitching artifacts with rdLight1 when stimulation and recording take place in the same brain region.^47^ To minimize this, we stimulated with comparatively low light power (5mW) in the SNc, >3mm away from the striatal recording site. Control stimulations in mice *not* expressing opsin indicated that rdLight1 recordings were not contaminated by blue light-induced photoswitching^47^ artifacts (“No Opsin” control mice, **Fig 3d; Extended Data Fig 5d**).

**Fig 3.**
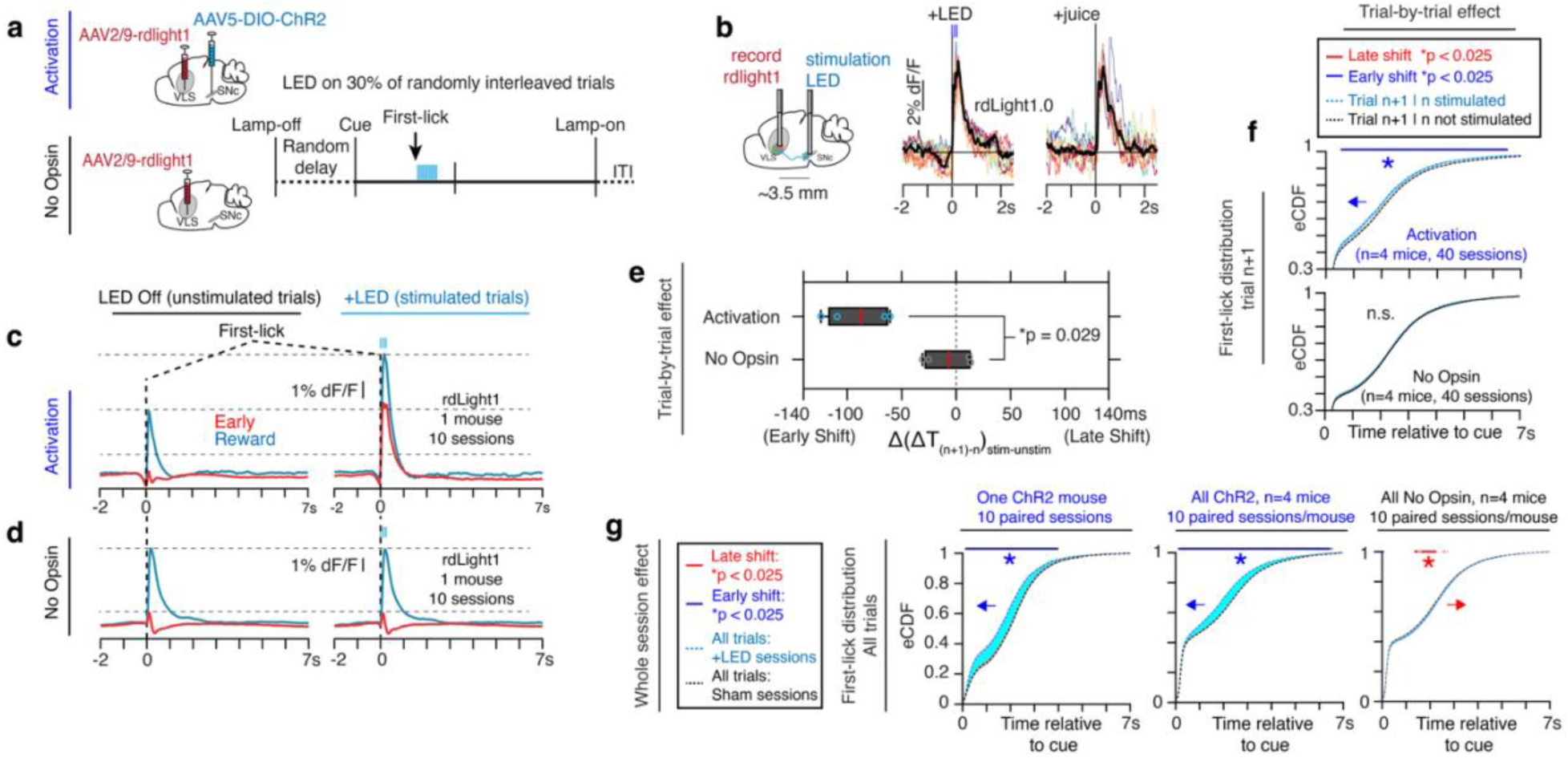
Outcome-related DA causes one-shot timing updates. a, Optogenetics strategy. **b,** Calibration of stimulation parameters outside the context of task. Mice received 10 unexpected juice rewards (∼4µL) followed by sets of 10 trials of optogenetic stimulation, with each set testing different stimulation parameters (Methods). Left: stimulation-aligned rdLight1 signals from trials using the optimal stimulation parameters in one calibration session. Right: juice-aligned signals from the same calibration session. Colored traces: 10x single trials, black traces: averages. **c-d,** Average VLS rdLight1 signals from all trials for one example ChR2 mouse (panel *c*) and one no opsin mouse (panel *d*). Slight differences in DA dynamics before the lick between panels *c-d* were within the normal range of inter-animal variability seen in this task.^7^ Data from all mice shown in Extended Data Fig 5. **e-f,** Single trial (“one-shot”) stimulation effects. **e,** Standard box plot. n=4 Activation mice; n=4 No Opsin mice, 10 sessions/mouse. Wilcoxon rank sum test, *p<0.05. **f,** First-lick empirical continuous distribution function (eCDF) on the next trial (n+1) after stimulated trials (blue trace) vs. after interleaved unstimulated trials (black trace) from the same session. Arrow: direction of effect. Line-shading: 95% CI; Blue bar above the plot indicates times where the eCDFs were significantly different (*p<0.025, 1000x bootstrap). **g,** Whole session effects: comparison of first-lick eCDFs of all trials from stimulation sessions (blue traces) vs. all trials from interleaved sham sessions (black traces). Area between the curves filled for clarity. Arrow: direction of effect. Line-shading: 95% CI; Blue/red bar above plots indicates times where eCDFs were significantly different (*p<0.025, 1000x bootstrap).

DAN activation caused mice to move significantly earlier than expected on the next trial, with 4/4 mice showing early-shifting (**Fig 3e-f**; see **Methods**; mean ΔΔT_(n+1)–n_:-88ms, p=0.0019, 10,000x bootstrap test, n=4 mice, 40 sessions). By contrast, stimulation in No Opsin mice had no significant effect on ΔT_(n+1)–n_ (mean ΔΔT_(n+1)–n_:-6ms, p=0.40, 10,000x bootstrap test, n=4 mice, 40 sessions). These results were consistent across 3 additional complementary statistical analyses (**Extended Data Fig 6**). Activation also caused early-shifting of the *entire* timing distribution across the session compared to interleaved sham sessions (**Fig 3g**; p<0.025, 1000x bootstrap test; **Extended Data Fig 5b,e**). This raised the possibility that outcome-related DA may normally control the “set-point” of the movement timing distribution, which we operationally define as the median first-lick time.

(We also attempted to optogenetically inhibit DANs at the time of reward, but similar to others (e.g, ref. ^32^ and Joshua Dudman, personal communication) we could not block the reward transient by triggering inhibition at the time of the first-lick within the context of the task, despite identifying stimulation parameters that caused DA dips outside the context of the task (n=4 stGtACR2 mice, see **Methods**). This was likely due to inability to initiate inhibition rapidly enough to prevent DAN bursting when light delivery was triggered on the first-lick. We thus examined reductions to outcome-related DA through non-optogenetic manipulations (discussed in the following sections).)

### DA explains the timing set-point

Surprisingly, across timing paradigms, both rodents and non-human primates frequently tend to move too early, at the expense of reward^7,40–43^ (**Extended Data Fig 1a**). In our mice, this early-bias developed gradually over days of experience (**Extended Data Fig 1c-e**) and persisted even in expert mice with >70 sessions’ experience.^7^ Outcome-related DA also appeared highly skewed in this task, with disproportionately large reward transients compared to small unrewarded dips (**Extended Data Fig 7a**). We observed that high amplitude (rewarded) DA transients caused much larger early-shifts compared to relatively small late-shifts after DA dips, and overall, mice made larger updates to timing after rewarded trials compared to unrewarded trials (median –317ms vs 74ms; **Fig 1g**). Together, this suggests bias in DA signaling may explain early-biased timing behavior.

But behavior was not early skewed for the entire session—all mice showed marked timing changes *within* each session, initially moving too early, but progressively shifting to later times (**Fig 4a-b; Extended Data Fig 7b**). If allowed to work *ad libitum*, the timing set-point (median of the timing distribution) ultimately shifted into the reward window—suggestive of timing calibration. (Importantly, the GLMs were robust to this non-stationarity, **Extended Data Fig 4c**).

**Figure 4.**
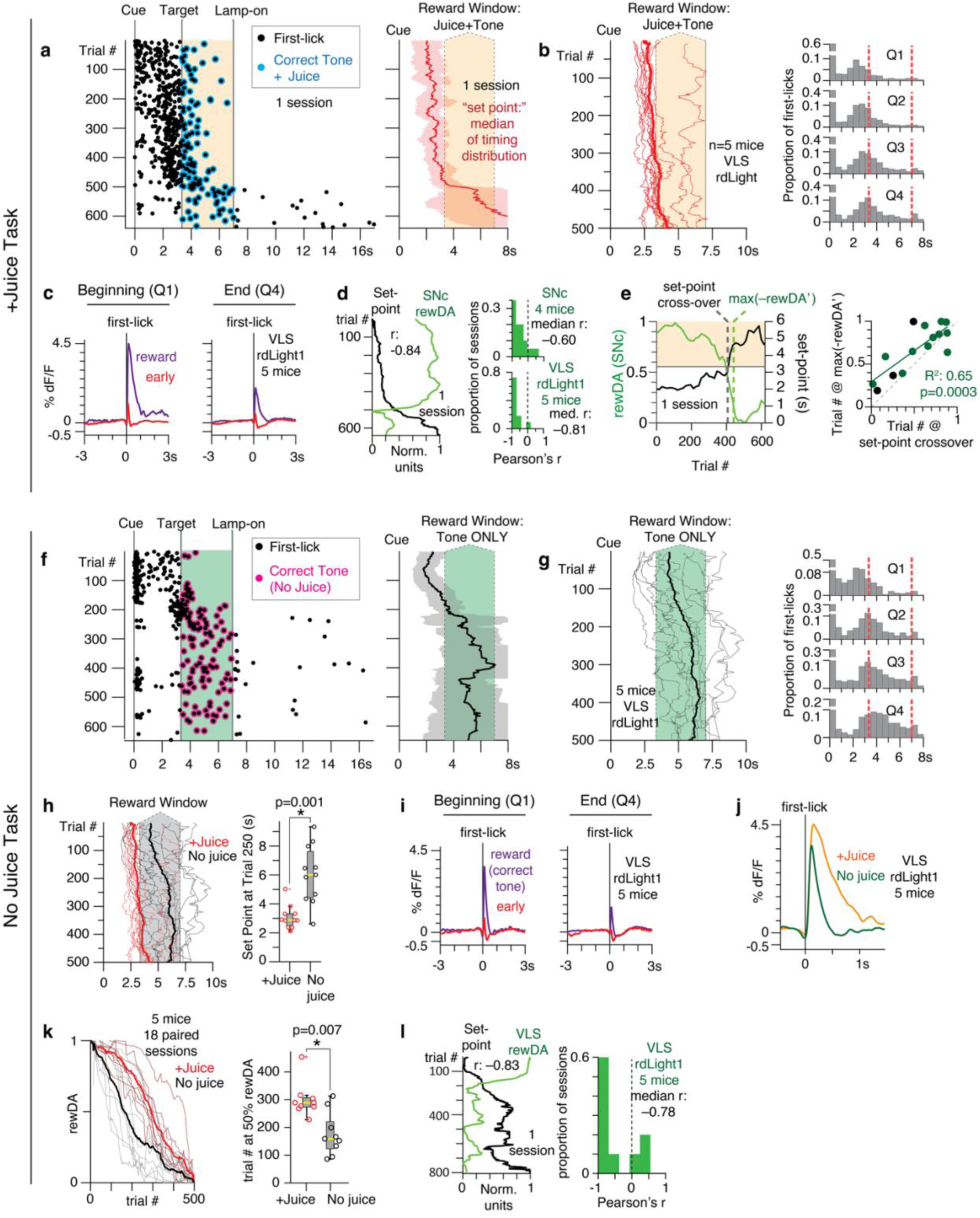
Timing behavior evolves over the course of each behavioral session. a, Trial-by-trial first-lick time across one entire session of the task. Left: raster of first-licks and rewards; Right: 50-trial moving median of first-lick times (shading: std). We define the *“set-point”* of the timing distribution on a given trial as the value of this moving median at that trial. **b,** Left: Set-point dynamics for different sessions and mice. 11 sessions from 5 mice are shown for comparison to data from the same mice on paired sessions in panels *f-l*. Right panel: Q1-Q4: distributions of pooled first-lick times across the same 11 sessions from the four quartiles of trials in each session. Timing distributions show gradual late-shifting over the course of the session. **c,** Average early and reward-related DA from the first (Q1) and last (Q4) quartiles of trials among sessions. **d,** Left: normalized peak rewarded DA transient amplitude (“rewDA”) vs. timing set-point for one example session. RewDA was defined as the maximum amplitude of the DA signal within 0.5s after the first-lick on single-trials, which was then interpolated for trial-by-trial comparison to the timing set-point. Right: distributions of correlation coefficients between rewDA and timing set-point for all SNc GCaMP6f and VLS rdLight1 sessions. **e,** Left: comparison of the time of the maximum negative derivative of rewDA (max(-rewDA’)) vs the trial at which the timing set-point first entered into the reward window (*“cross-over” point*; one example session). Right: max(-rewDA’) vs cross-over point for all individual SNc GCaMP6f and VLS rdLight1 sessions that exhibited a cross-over of the timing set-point into the reward window. Green points: +Juice sessions. Black points: No Juice task sessions. **f-g:** No Juice Task; as in panels *a-b*. Panel *g* shows all 11 paired No Juice sessions from the same mice as in panel *b*. **h,** Comparison of timing set-points for all paired +Juice vs No Juice sessions. Boxplot: distributions across sessions of the position of the timing set-point measured at trial 250; yellow lines: medians of the distributions; p: Wilcoxon rank sum. **i,** No Juice Task outcome-related DA. **j-k:** Comparison of rewDA in the +Juice vs. No Juice Task. Panel *j* shows Q1 signals from each task; Panel k’s boxplot shows the distribution of trial-number at which rewDA reached 50% of its max value; p: Wilcoxon rank sum. **l,** No Juice rewDA vs timing set-point, as in panel *d*.

We hypothesized that changes in outcome-related DA could explain changes in the timing set-point. Across each session, as the set-point shifted later, 1) rewarded trials were followed by progressively smaller timing updates (median timing change (in excess of chance), **beginning half of session:-400ms**, 95% CI: [-519ms,-270ms]; **ending:-271ms**, 95% CI: [-403ms,-143ms]; n=17 mice, 155 sessions; **Extended Data Fig 7c**); and, 2) rewarded DA transient amplitudes progressively declined (**Fig 4c; Extended Data Fig 7d-e, Extended Data Fig 8**). Importantly, this DA decline was not explained-away by bleaching or optical artifacts (**Extended Data Fig 7f**, see **Methods**).

Rewarded DA amplitude was tightly correlated with the timing set-point (median Pearson’s r VLS rdLight1: –0.81; SNc GCaMP6f: –0.60; **Fig 4d**). This was not the case for movement-related/optical artifacts (median Pearson’s r tdTomato: +0.19, **Extended Data Fig 7g**). Strikingly, shifts were not always gradual—rewarded DA amplitude decreased suddenly during abrupt shifts of the set-point into the reward window (**Fig 4e**; R^2^=0.65, p=0.0003), which was also not explained away by optical/movement related artifacts (R^2^=0.19, p=0.10; **Extended Data Fig 7h**).

### DA calibrates the propensity to move

Altogether, across sessions and recording sites/modalities, rewarded DA transients decreased in amplitude as timing behavior shifted later, toward the reward window. DA transients also controlled trial-by-trial timing updates, and these updates became progressively smaller as behavior became more accurate (**Extended Data Fig 7c**). Together, these findings suggest a calibration process for motor timing. This adaptive decrease in stochastic learning is reminiscent of adaptive stochastic gradient descent algorithms.^48^ However, because DA-reward transient amplitude and set-point tended to change monotonically across the session, we tested several complementary manipulations to “break the monotonicity” between these co-variates. Each manipulation produced marked changes in the animals’ timing strategy, including late-to-early timing shifts. Importantly, in all settings, DA reward transients explained bidirectional changes in the timing set-point, as follows:

### Different reward identities

First, we serendipitously discovered that well-trained mice continued to participate even when juice rewards were replaced by the usual “correct” tone (**“No Juice” Task**; solenoid valve disconnected)—but set-point dynamics were strikingly different (**Fig 4f-g**; **Extended Data Fig 7i**). Mice still showed early bias for the first ∼150 trials, but unexpectedly, the set-point suddenly and dramatically shifted into the reward window—significantly earlier in the session than in the +Juice task (p=0.001, Wilcoxon rank sum test; 5 mice, 22 paired behavior sessions, **Fig 4h**). Mice continued to work for as many as 1000 trials under these conditions with correct timing on most trials, exhibiting much more “accurate” behavior than the same animals performing the +Juice task on interleaved sessions (same mice as **Figs 1b, 4b-d**).

Surprisingly, mice still showed DA “reward” transients in the No Juice task (in response to the correct tone, **Fig 4i**), but these were narrower and smaller in amplitude than reward transients in the +Juice task (**Fig 4j**). Moreover, reward-transient amplitude declined significantly earlier in the session during No Juice sessions (n=5 mice, 18 paired photometry sessions; p=0.0073, Wilcoxon rank sum test, **Fig 4k, Extended Data Fig 7j-l**), and the timing of this abrupt, early decline explained the rapid shift in timing set-point (median Pearson’s r: –0.78; **Fig 4l; Extended Data Fig 7n**). These effects were not explained by changes in ongoing body movements or optical artifacts (**Extended Data Fig 7k,m**).

Interestingly, the persistence of high amplitude reward transients in the +Juice task suggests that juice skewed normal timing updates: despite apparent motivation, mice received many fewer rewards/time compared to the No Juice task. The resulting early timing bias is reminiscent of the effects of DA-modulating drugs of abuse on timing behavior, such as amphetamines.^37^

### Abrupt changes in reward identity

Second, we trained mice to alternate between blocks of trials with either No Juice rewards (correct tone only) or +Juice. During No Juice blocks, the timing set-point rapidly late-shifted into the reward window. This was rapidly reversed in +Juice blocks, in which the timing set-point rapidly shifted significantly earlier (n=3 mice, 5 block changes; p=0.032, Wilcoxon rank sum test; **Fig 5a**). Again, the amplitude of DA reward transients explained these set-point shifts (median Pearson’s r: –0.89; **Fig 5b-c**).

**Figure 5.**
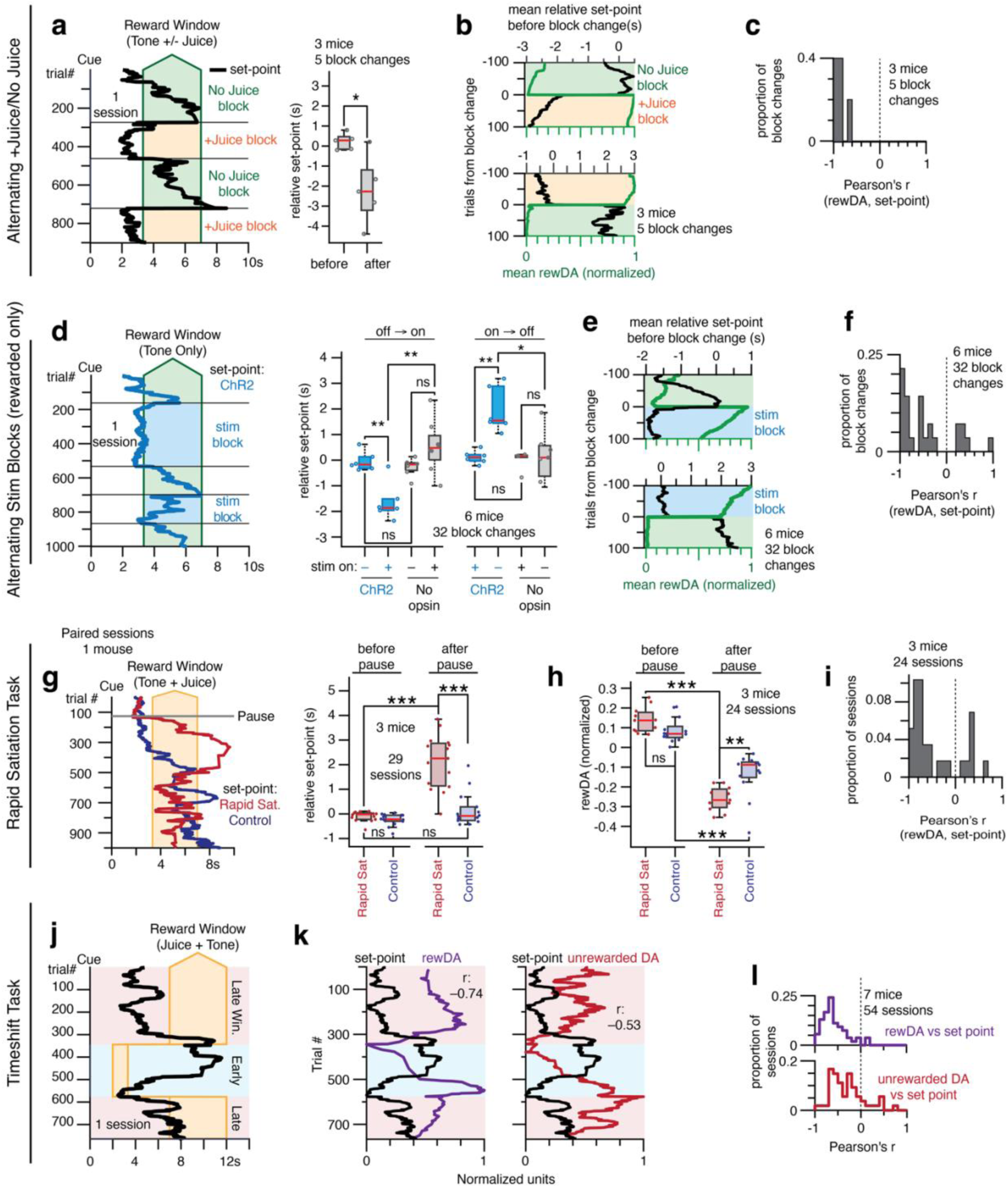
Outcome-related DA transients calibrate the propensity to move. a-c, Left: timing set-point in the Alternating Juice/No Juice Task, with set-point defined as the 35-trial moving median of the first-lick time distribution, as in Fig 4 (see Methods). One example session shown. Right: comparison of timing set-point before and after all block changes from all mice, all sessions at transitions from No Juice to +Juice blocks, where each block change’s relative set-point was calculated as the average set-point over the 100 trials before or after the block change minus the set-point measured on the last trial before block change (negative number indicates early-shift of timing set-point relative to the set-point at the block change). Standard box plot, Wilcoxon rank sum, p*=0.032. **b-c,** Correlation of reward-related DA (“rewDA,” as defined in Fig 4) and timing set-point across block changes. Left: average set-point and rewDA (all block changes, all mice); Right: correlation coefficient of set-point and rewDA for each block change (all block changes, all mice). **d-f,** Alternating Stimulation Blocks, Rewarded Trials only. **d,** Left: timing set-point for one example stimulation session. Right: comparison of set-point across block changes as in panel *a* (ChR2 vs No Opsin animals, all block changes, all mice). *p<0.05, **p<0.01, ***p<0.001; Wilcoxon rank sum. **e-f,** Correlation of set-point and rewDA across block changes, as in panels *b-c*. **g-i,** Rapid Satiation task. **g,** Left: Comparison of timing set-point in a paired rapid satiation session (red trace) vs. a control session (purple trace) from the same mouse. “Pause” indicates trial at which mouse was given free access to juice (rapid satiation session) or an empty syringe (control). Right: comparison of timing set-point before and after the pause (all mice, all sessions), as in panel *d*. Positive relative set-point indicates a late-shift of the set-point compared to the set-point on the last trial before the pause. **h**, Comparison of rewDA before and after the pause (all photometry sessions, all mice). RewDA amplitudes for each session normalized to rewDA on the last trial before the block change, such that a negative value indicates a decrease in rewDA. **j-l,** Timeshift Task. **j,** Timing set-point (one example +Juice session). **k,** Comparison of set-point and rewDA (left panel, as in Fig 4d) or unrewarded DA (right panel, max DA signal within 500ms of first-lick on unrewarded trials, including both early and late trials). One example session. **l,** Distribution of correlation coefficients for set-point and rewDA (purple) or unrewarded DA (red) across all sessions and mice. Set-point remains strongly correlated with rewDA; unrewarded DA peak amplitude shows stronger correlation with set-point than in the standard +Juice/No Juice tasks.

The rapid changes in DA-transient amplitude might be correlated—but not causal—to abrupt set-point shifts. Therefore, we also had mice alternate between blocks of trials with either No Juice or No Juice+Stimulation, in which DANs were optogenetically activated at the time of first-licks that occurred either within the reward window or on all trials (n=3 mice, 20 block changes; **Extended Data Fig 9a-e**). Stimulation was calibrated to recapitulate usual juice-related DA transients without delivering liquid rewards (as in **Fig 3b**). The set-point rapidly late-shifted during No Juice blocks, but then rapidly and significantly early-shifted during +Stimulation blocks (off-on: p=0.001, on-off: p=0.007; Wilcoxon rank sum test; **Extended Data Fig 9b**), even when stimulation was only delivered on rewarded trials (off-on: p=0.009, on-off: p=0.002; n=2 mice, 12 block changes, **Fig 5d**). Stimulation did not change the set-point in No Opsin control mice (n=4 mice, 20 block changes; **Fig 5d; Extended Data Fig 9b-e**). Again, the amplitude of DA reward transients explained the set-point shifts (median Pearson’s r, stim: –0.66, no opsin: –0.56; **Fig 5e-f, Extended Data Fig 9d-f**). These results emphasize that DA-reward transients induced *shifts* to earlier times—rather than reinforcing the rewarded movement time.

### Rapid satiation

Third, in addition to sudden shifts, the timing set-point also showed gradual late-drift across sessions, potentially related to fatigue or satiation. We hypothesized that DA could also explain state/motivation-dependent shifts. We thus tested whether *rapid* satiation could change DA reward transients and set-point. After completing >=100 trials of the +Juice task and receiving >=10 rewards, the task was paused, and mice were allowed to drink *ad libitum* until they stopped licking available juice (n=3 mice, 15 sessions; range: 0.75-1.8mL (daily water consumption for a water-deprived mouse is ∼1mL); **Fig 5g-h**). The task was then resumed. DA-reward transient amplitude and timing set-point remained highly anticorrelated throughout the session (median Pearson’s r: –0.70, **Fig 5i**), but DA-reward transient amplitudes decreased rapidly after the pause (p=8.5e-5, Wilcoxon rank sum test; n=11 photometry sessions; **Extended Data Fig 9g**), and the timing set-point showed significant late-shifting (p=1.7e-5, Wilcoxon rank sum test; **Extended Data Fig 9h**). By contrast, in interleaved control sessions (empty syringe; no juice given during the pause), rewarded DA did not show this precipitous decrease (p=0.0026; **Fig 5h**), and there was no significant change in timing set-point (p=0.18; **Fig 5g**). Altogether, rapid juice consumption caused subsequent DA reward transients to rapidly decrease in amplitude, which was associated with late-shifting.

### Changing timing constraints

Finally, fourth, to test if endogenous changes in DA reward transients could re-calibrate the timing set-point in response to different environmental timing contingencies, we changed the rewarded time window in blocks of trials (“Timeshift Task;” e.g., from 7-12s to 2-3.3s; n=7 mice; **Fig 5j**). The new reward window was not signaled and had to be discovered through trial-and-error. We hypothesized increases in DA-reward transient amplitdues would cause early-shifting to match the constraints of the task (and decreases, late-shifting). Because fatigue/satiety should increase monotonically over the course of the session, we reasoned that early-shifts would disentangle goal-directed processes from fatigue/satiation.

The Timeshift Task produced pronounced shifts in the animals’ timing behavior, including directed changes in timing in relation to the new reward window (**Fig 5j, Extended Data Fig 9i**) and exploratory behavior that appeared to anticipate block changes (**Extended Data Fig 9j**). The amplitude of rewarded DA transients continued to explain the timing set-point (median Pearson’s r: –0.64; 7 mice, 54 sessions; **Fig 5k-l**), and intriguingly, *unrewarded* DA transients also became more correlated with the set-point (median r: –0.36; **Fig 5l**). Early-shifts were accompanied by increases in DA-transient amplitude (and late-shifts by decreases), and these effects were not explained away by optical artifacts (tdt: Rewarded trials median r: –0.20; unrewarded r: 0.10; **Extended Data Fig 9k**).

Together, these results show that changes in reward-related DA transients cause shifts in the behavioral timing set-point, both in relation to changing environmental conditions (such as changes in environmental timing contingencies for reward or the presence/absence of juice rewards) and internal state (such as gradual or rapid satiation). These findings demonstrate that DA reward transients quantitatively and adaptively calibrate the propensity to move in response to both external and internal constraints.

## DISCUSSION

A cornerstone hypothesis of biological reinforcement learning is that reward-related DA transients (whatever their nature^3,5,16,19,49,50^) lead to changes in future behavior. However, to our knowledge, variability in endogenous reward-related DA transients has not been shown to relate to variability in subsequent behavior, despite serious effort to detect such effects.^3,5,31,51,52^ Here, we show for the first time that the amplitude of native DA reward transients is quantitatively related to one-shot, goal-directed updates to movement timing on the *next* trial, >10s later. This suggests a functional link between reward and the calibration of the propensity to move (an aspect of vigor), unifying two seemingly distinct aspects of DA function.^6^ In addition, we found that asymmetry in the relative amplitude of rewarded/unrewarded DA transients explained asymmetry in one-shot updates, and that changes in the relative amplitude of these transients could causally shift the timing set-point. This process, reminiscent of adaptive stochastic gradient descent,^53^ allowed the animals to adjust their propensity to move in response to changing experimental contingencies (e.g., different timing constraints). Changes in reward-related DA transients and behavior could also be induced by changes in internal/motivational state (reward identity, satiety/fatigue), demonstrating that flexibility in reward-related DA signaling mediates both strategic and motivationally driven changes to movement timing. Together, DA’s tripartite roles in reward, movement and motivation are mechanistically linked via this calibration process.

### Relationship to previous results

We and others have observed “real-time” effects of DA on movement—but over much shorter timescales than the reward-related effects we report here (within 100s of milliseconds before or during ongoing movements).^7,23–26^ Others have suggested that DA’s endogenous effects on vigor are instead related to slowly-evolving or “tonic” DA signals rather than trial-by-trial reward-related DA transients.^6,22,27,29^ Here we found that variability in DAN baseline activity (before the start-timing cue) could be traced back *tens of seconds* to variability in the amplitude of the previous trial’s outcome-related DA transient—and that variability in this transient can causally update timing on the next trial, 10-25s in the future, perhaps by establishing a putative “stochastic set-point” that controls the start-point and slope of the ramping process.^54–56^

*Exogenous* DA manipulations also affect future behavior, but typically only gradually, over many exposures.^10,12,14,57,*c.f.*32^ Endogenous DA-transient amplitude can also decrease over the course of learning in a variety of settings,^2,22,58,59^ but again only gradually, and also not linked to changes in behavioral vigor.^22^ Other recent work showed an increased repetition of behavioral motifs that had been associated with spontaneous DA transients in freely behaving mice,^11^ but this was outside a goal-directed context and did not explore the effects of rewards or reward-related DA transients.

Why could we detect one-shot effects of endogenous DA-reward transients on behavioral vigor when previous studies did not? One possibility is that our time-to-move metric provided a more sensitive “analog” behavioral readout. Behavioral updates in discrete reinforcement-learning (RL) tasks, such as bandit tasks, may be difficult to detect on a trial-by-trial basis,^3,51,57^ instead requiring multiple repetitions before becoming manifest. Other work examining behavioral vigor in freely behaving animals (measured by the animals’ latency to initiate trials) may have missed one-shot effects of DA-reward transients due to high variance and long latency between trial-starts, as long as 30s.^4,5^ The one-shot effects we observed—which were on the order of tens to hundreds of milliseconds—would be difficult to detect in this setting. However, we also found that DA’s one-shot effects accumulated over trials, which could potentially account for the correlation between DA signals and vigor effects when averaged over long time periods.^4,5^

Interestingly, a recent mouse knockout model of action potential-driven DA transients (RIM^cKO^) caused surprisingly few gross motor deficits in these animals, although non-phasic DA release was still necessary for movement.^27^ A priori, we might expect the trial-by-trial timing updates that we observed would be absent in these mice, potentially preventing the animals from flexibly calibrating their timing based on trial outcome.

### Proposed model

DA signals are often interpreted in the framework of normative RL models. In the simplest model, rewards might reinforce rewarded lick times, but we found that DANs instead drove *changes* in timing on the next trial. This could still be consistent with RL algorithms that learn to anticipate reward-relevant events, such as temporal difference (TD) learning^60^ (and others). If so, an informative case would be the infrequent *late* trials in our task, where animals first licked after the end of the reward window and thus were not rewarded. RL algorithms for updating behavioral policies, such as Q-learning,^61^ could, in principle, learn to promote earlier movements after late trials to increase the probability of reward. But instead, we found that late trials led to *late*-shifting on the next trial (**Fig 1g**), in conjunction with DA dips (**Extended Data Fig 10a-b**). Although extensions of RL models might be able to explain this effect, any such model would run counter to the normative goal of maximizing future rewards—which is the premise of RL algorithms. Parsimony thus suggests a different model than RL. We propose that DA transients rapidly—and in a *fixed manner irrespective of trial outcome*—modulate the propensity to move *proportional to the amount of DA released* (**Extended Data Fig 10c-e**), with larger DA transients leading to increased vigor (earlier movements) and smaller transients, such as dips, leading to decreased vigor (later movements). In this view, rapid one-shot DA effects may not be related to “learning” *per se*, but could reflect an “innate,” automatic mechanism for quickly modulating vigor in response to action outcomes. This rapid mechanism could explain how DA-modulating drugs, such as amphetamines, increase vigor automatically and non-specifically without apparent learning.^62^ Consistent with this idea, we found a similar negative correlation between reward-related DA transients and trial-by-trial changes in the timing of movement when we re-analyzed data from an unrelated cued reaching task,^63^ in which the timing of reaches had no bearing on whether reward was delivered or omitted (**Extended Data Fig 10f-h**).

Importantly, we are not arguing against DA’s role in RL. On the contrary, rapid DA effects on vigor could operate in parallel—or even be synergistic—with slower, incremental DA-mediated RL. For example, rapid vigor modulation may have evolved to promote movement in reward-rich environments, which would increase the number of opportunities to repeat rewarded actions, which in turn could drive more gradual, persistent and selective RL. This interpretation could explain DA’s association with exploratory behavior,^6^ as well as the observation that animals increase the frequency and speed of all movements, not just reward-associated movements, in reward-rich environments.^6^

### Potential biological mechanisms

Theoretical work has suggested frameworks for understanding DA’s effects on vigor, but the biological implementations were not clear.^6,64^ We previously showed that physiologically calibrated optogenetic DAN activation (or inhibition) during the self-timed interval rapidly (sub-second to a few seconds) increased or decreased the moment-by-moment probability of initiating the timed movement.^7^ Those rapid effects were in the same direction as the effects of the outcome-related DA transients in the current study (i.e., higher DA leads to earlier movement), presumably reflecting a common mechanism. Therefore, any proposed biological mechanism should have the following properties: 1) it should support *bidirectional* effects on timing; 2) it should have a *rapid onset* (sub-second to a few seconds); and 3) it should *persist* for at least tens of seconds (the duration of the ITI in our experiments).

Gradual DA-dependent structural synaptic plasticity in cortico-striatal synapses has been proposed as a biological mechanism by which DA could mediate learning.^65^ However, synaptic plasticity is likely too slow and incremental to explain the rapid effects on behavioral vigor that we observed—and would also seem metabolically inefficient to support back-and-forth, trial-by-trial updates. We thus suspect that one-shot effects on vigor arise via a fundamentally different biological mechanism than synaptic plasticity.

DA also modulates the excitability of striatal projection neurons (SPNs) via metabotropic receptors.^66,67^ A salient example is a careful striatal slice study,^67^ which found that calibrated, transient optogenetic stimulation of DAN axons caused rapid (sub-second) yet long-lasting (minutes) increases in the firing rate of direct pathway dSPNs in response to depolarizing current injections.^67^ This excitability effect was blocked by a specific inhibitor of Protein Kinase A (PKA). PKA is an intriguing candidate effector, because DA transients *in vivo* have been shown to drive increased PKA activity in dSPNs, whereas DA dips cause increased PKA activity in indirect pathway iSPNs, with a latency of a few seconds and persistence for tens of seconds,^68^ similar to the dynamics of the behavioral effects we observe. It is possible that the relative balance of dSPN:iSPN excitability could scale dynamics of the cortico-basal ganglia-thalamic circuit^62,69^ in a manner that allows for bidirectional changes in timing—and modulation of vigor more broadly. Outcome-related DA transients could drive these PKA changes. This scaled-dynamics view provides a framework for understanding vigor-related symptoms of DA-related disorders of movement, such as bradykinesia and akinesia in Parkinson’s disease.

## METHODS

### Animals

We used adult male and female hemizygous DAT-cre mice^70^ (B6.SJL-*Slc6a3^tm1^*^.1(cre)Bkmm^/J, RRID:IMSR_JAX:020080; The Jackson Laboratory, ME) or wild-type C57BL/6 mice (>2mo old). Mice were housed in a temperature and humidity-controlled facility on a reversed night/day cycle (12h dark/12h light). Behavioral sessions occurred during the dark cycle. All protocols were approved by the Harvard Institutional Animal Care and Use Committee and MIT Committee on Animal Care (Harvard IACUC protocol #1343, Animal Welfare Assurance Number #A3431-01; MIT CAC protocol #2306000543). All work was in accordance with the NIH Guide for the Care and Use of Laboratory Animals. **Surgeries** were conducted under aseptic conditions with every effort to minimize suffering, as we described previously.^7^ Mice were **water deprived** as we described previously^7^ (≥80% initial body weight) with *ad libitum* feeding. **Histology** to confirm viral expression and fiber placement was conducted as we reported previously.^7^

### Stereotaxic coordinates (from bregma and brain surface)

Viral Injection:

SNc^7^: 3.16mm posterior, +/-1.4mm lateral, 4.2mm ventral VTA^7^: 3.1mm posterior, +/-0.6mm lateral, 4.2mm ventral DLS^7^: 0mm anterior, +/-2.6mm lateral, 2.5mm ventral.

VLS^45^: 0.34mm anterior, +/-2.75mm lateral, 3.7mm ventral (rdLight1/dLight1.3), 3.2mm ventral (D1/A2A-cre; GCaMP)

Fiber Optic Tips (ventral position): 200µm 0.53NA or 0.55NA blunt fiber optic cannulae (Doric Lenses, Quebec, Canada) or tapered fiber optic cannulae (200 µm, 0.60 NA, 1.5, 2 or 2.5mm tapered shank, OptogeniX, Lecce, Italy) were positioned at SNc, VTA, DLS or VLS: SNc/VTA^7^: 4.0mm (photometry) or 3.9mm (optogenetics).

DLS^7^: 2.311mm (blunt) or 4.0mm (tapered) VLS (rdLight1/dLight1.3): 3.4mm (blunt)

VLS^45^ (D1/A2A-cre; GCaMP): 3.0mm (blunt or midpoint of 2.5mm tapered)

### Viruses

Photometry:

<u>Optical control fluorophores</u>: 500nL alone or mixed with other fluorophores (below)

*-tdTomato (“<u>tdt</u>”)*: AAV1-CAG-FLEX-tdT (5.3*10^10^ to 5.3*10^12^gc/mL; UNC Vector Core, NC)
*-<u>GFP</u>*: AAV9.CB7.CI.eGFP.WPRE.rBG (2.3*10^12^ vg/mL; Addgene plasmid #105542)
<u>gCaMP6f (at SNc or VTA)</u>: 300nL AAV1.Syn.Flex.GCaMP6f.WPRE.SV40 (2.5*10^13^gc/mL;

Penn Vector Core, PA) mixed 3:1 with tdt or mCherry (D1/A2A-cre experiments; AAV8-Syn-DIO-mCherry, 2*10^12^gc/mL; Datta Lab)

DA_2m_ (at DLS): 200-300nL AAV9-hSyn-DA4.4(DA2m) (3*\**10^12^gc/mL; Vigene, MD) mixed 2:1 or 3:1 with tdt

dLight1.1 (at DLS): 300nL AAV9.hSyn.dLight1.1.wPRE (9.6*10^12^gc/mL, Children’s Hospital Boston, MA) mixed 3:1 with AAV1.CB7.CI.TurboRFP.WPRE.rBG (10^12^gc/mL, Penn Vector Core)

rdLight1 (at VLS): 500nL AAV2/9-Syn-RdLight1.0 (2*10^12^gc/mL; #construct-1646-aav2-9, Canadian Neurophotonics Platform (CNP), Quebec, Canada).

dLight1.3 (at VLS): 500nL AAV2/DJ-CAG-dLight1.3b (2*10^12^gc/mL, CNP) Optogenetic stimulation/inhibition (bilateral at SNc):

ChR2: 1000nL AAV5-EF1a-DIO-hChR2(H134R)-EYFP-WPRE-pA (3.2*10^13^gc/mL, UNC Vector Core, NC)

<u>stGtACR2</u>: 300nL AAV2/8-hSyn1-SIO-stGtACR2-FusionRed (4.7*10^11^gc/mL, Addgene/Janelia Viral Core, VA)

### Behavior

#### Self-timed movement task (original “+Juice” version)

Mice were head-fixed with a juice tube positioned in front of the tongue. The spout was placed ∼1-1.5mm ventral and ∼1-1.5mm anterior to the mouth. During periods when rewards were not available, a houselamp was illuminated. At trial start, the houselamp turned off, and a random delay ensued (0.4-1.5s) before a cue (simultaneous LED flash and 3300Hz tone, 100ms) indicated start of the timing interval. The timing interval was divided into two windows, early (0-3.333s) and reward (3.333-7s), followed by the intertrial interval (ITI, 7-17s). The window in which the mouse first licked determined the trial outcome (early, reward, late or no-lick). An early first-lick caused an error tone (440Hz, 200ms) and houselamp illumination, and the mouse had to wait until the full trial interval had elapsed before beginning the ITI. Thus, there was no advantage to licking early. A first-lick during the reward window caused a reward tone (5050Hz, 200ms) and juice delivery (∼4µL apple juice or Gatorade; **+Juice Task**), and the houselamp remained off until the end of the trial. If the timing interval elapsed with no lick, a time-out miss tone played (131Hz, 2s), the houselamp turned on, and ITI commenced. During the ITI and pre-cue delay (“lamp-off interval”), there was no penalty or feedback for licking. Mice learned the task in 3 stages (beginner, intermediate and expert), as described previously.^7^ All mice learned the task and worked for ∼400-1,800 trials/session.

#### Pre-training/post-training

To encourage participation, each session (except for DAN GCaMP mice) commenced with 10 trials of “pre-training” during which juice was delivered triggered on the first-lick within the reward window, but during which early licks were not penalized. Pretraining was always conducted with juice rewards.

#### Quality control standards

Rare trials where the first-contact with the spout was a paw contact rather than a lick (e.g., due to grooming) and rare trials with rig software/hardware errors (e.g., clogging of the juice spout) were excluded from analysis. Analyses were robust to inclusion/exclusion of these trials. We provide all exclusions and raw data in the public datasets so that users can decide whether to include these or not.

#### One-shot Stimulation (Fig 3)

Closed-loop stimulation was delivered at SNc and triggered on the first-lick on 30% of randomly interleaved trials in the context of the +Juice Task. Mice were trained without stimulation for 13-16 sessions before starting stimulation sessions. Each mouse completed 10 pairs (20 sessions) of interleaved stimulation and sham sessions, where sham sessions were identical to stim sessions (patch cables attached, etc.), except the stimulation LED was turned off and no light was delivered. Photometry was usually not collected during sham sessions to reduce bleaching from recording on consecutive days. See **Optogenetics** section for stimulation parameters, calibration and analysis.

#### No Juice Task (Fig 4f-l)

In the usual (+Juice) task, feedback was either an error tone (440Hz, 200ms) or correct tone (5050Hz, 200ms) plus juice reward. The No Juice Task was identical to the +Juice Task and used the same feedback tones, but no juice was delivered (dispensing solenoid valve disconnected). In a subset of animals trained on an open rig with a loud juice solenoid, a decoy solenoid was triggered, but no juice was dispensed. After >=35 days of +Juice Task, expert mice successfully performed the No Juice Task, some for >1000 trials/session (one mouse tested on days 23 and 25 showed less engagement). Anecdotally, we observed that consecutive No Juice sessions could reduce participation in subsequent sessions in some animals, and so No Juice sessions were typically interleaved with +Juice sessions to promote participation. Together, these factors limited the number of No Juice sessions that we could collect.

#### Alternating Juice/No Juice Task (Fig 5a-c)

Mice alternated between unsignaled blocks of the +Juice or No Juice Task.

#### Alternating Stimulation Blocks (Fig 5d-f)

Mice were trained in a No Juice Task variant in which closed-loop SNc DAN stimulation was delivered in blocks of trials. During stimulation blocks, stimulation was triggered on either 30%, 60% or 100% of first-licks (**Extended Data Fig 9a**) *or* only on 100% of first-licks occurring within the reward window (**Fig 5d**). See **Optogenetics** section for stimulation parameters, calibration and analysis.

#### Rapid Satiation Task (Fig 5g-i)

At >20d of training, Rapid Satiation sessions were collected, where the ongoing task was paused once behavior met two criteria: ≥100 trials completed and ≥10 juice rewards received. Video and photometry recordings were continued during the pause (mice remained headfixed on the behavior rig). The spout was retracted, and the mice were hand-fed Cool Blue Gatorade from a 1mL syringe until they stopped initiating licks to the offered syringe. The spout was then moved back into position and the task resumed. In Control sessions, the same procedure was followed, except an empty syringe was offered.

Both Rapid Satiation and Control sessions were collected on a background of the +Juice task with Cool Blue Gatorade rewards. We used a newer version of the +Juice task, in which there was also an unsignaled late window after the reward window (7-14s). In this version of the task, the trial duration was always 14s, with the late window occurring between the unsignaled end of the reward window and the start of a 3s-long ITI (total trial+ITI duration 17s, as in the original +Juice Task). First-licks during the late window triggered a distinct, late error tone (132Hz, 500ms) and illumination of the houselamp (similar to early trials). Mice still had to wait until 14s to start the ITI.

In analyses, to account for missing time while the task was paused, the total duration of the pause was divided by 17 (the trial length was 17s) and rounded to recover the number of missing trials. Rewarded DA transient amplitudes were then interpolated linearly and smoothed with a 20-trial gaussian kernel (“rewDA”). Edges (trials before the first reward and after the last reward in the session) were omitted.

Rapid Satiation sessions were collected in pairs with interleaved control sessions. However, some sessions had to be excluded either because the mouse received no rewards before the pause (1 session), the mouse did not resume participation after the pause (thus precluding measurement of reward transients, 1 session), or there was failure of lick detection during the session due to a faulty spout (3 sessions). Although including these sessions did not change the result of analyses, they were omitted from the final rewDA analysis. Statistical comparisons of rewDA before and after the pause were performed on the average rewDA signal across 50 trials before and after the maximum duration pause (36 trials) and compared by Wilcoxon rank sum test.

#### Timeshift Task (Fig 5j-l)

In the usual +Juice and No Juice tasks, there was only one rewarded time window (3.333-7s). In the Timeshift Task, this window changed dynamically in blocks of trials. The Timeshift Task was introduced after >28 days of training on the usual +Juice Task. Each block used one of five reward windows: a) 1-2s, b) 2-3.333s, c) 3.333-7s, d) 7-12s, e) 9-14s. There was no set schedule for changing between blocks. Block changes were signaled by a ∼90s timeout (task paused). The new reward window was not indicated and had to be discovered through trial-and-error. In a subset of sessions, a “late” window was added (as in the Rapid Satiation Task) such that mice were not immediately alerted when the reward window had closed. Behavior was grossly similar on both versions of the Timeshift Task, and thus, for all analyses, data from the standard Timeshift Task and late window Timeshift Task were pooled. Mice were also trained on a No Juice version in some Timeshift sessions (7 mice, 9 sessions), and these data were pooled with +Juice Timeshift sessions.

### Regression to the median analyses

Timing behavior may arise from a stochastic generative process, in which timing on any trial can be modeled as stochastic “draws” from a latent distribution, *T*. This presents potential confounds to detecting trial-by-trial learning. In this framework, the expected median of the next trial’s movement time would be the median of the latent distribution on the next trial, *T^(n+^*^1^*^)^*. If the latent distribution is stationary trial-by-trial (i.e., *T^(n)^* = *T^(n+1)^*), we would still see changes in movement timing on the next trial (Δt_n_) that arise by chance, but which are not necessarily in response to learning. Thus, even though some deterministic process (say, outcome-related DA signaling) might control this tendency to “regress to the median” of *T^(n)^*, we could not rule out the possibility that regression to the median of *T^(n)^* happens by chance (i.e., due to a different process). Thus, to detect learning effects in excess of chance would require that Δt_n_ be in excess of the regression to the median effect. In a stochastic generative model of timing, we would interpret this kind of learning as an update to the latent distribution on the next trial, such that *T^(n)^* ≠ *T^(n+1)^.* We define regression to the median on each trial, n, as:

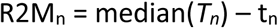

Where t_n_ is the movement time on trial n and *T_n_*is the vector of all movement times in consideration. We say “in consideration” because we cannot simply use all movement times in the session due to slow, non-stationarity in the timing distribution across the session, which might arise from processes other than learning (e.g., satiation/fatigue) and which necessitates a more local calculation of median(*T_n_*) to isolate local non-stationarity arising from trial-by-trial updates. Thus, the median(*T_n_*) is an uncertain quantity. To address this, we calculated median(*T_n_*) in 3 complementary ways:

A. We broke each session into deciles of trials and calculated the median of the distribution within each decile.
B. We broke the session into deciles as in A, but we bootstrapped the next trial’s movement time by resampling the distribution of movement times within the decile 10,000 times.
C. We calculated a local, moving median movement time for each trial (100 trials, centered at current trial).

R2M_n_ for each method was then compared to the true Δt_n_ on each trial, defined as in the rest of the paper (Δt_n_ = t_n+1_– t_n_; where late-shifting is positive by convention). Because of the high level of trial-by-trial variability, we resolved Δt in excess of R2M as a significant difference between the distributions of R2M_n_ vs Δt_n_ across all trials (R2M and ΔT), calculated by Wilcoxon rank sum test. The median of each distribution was then used to calculate “ΔT in excess of chance” (**Fig 1g**) defined as:

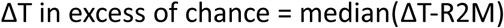

Assuming timing arises by a stochastic generative process, this excess term is interpreted as the change in the median of *T^(n+1)^* versus *T^(n)^*.

For the analysis dataset, we used all +Juice photometry sessions that recorded DAN/DA signals from mice expressing no opsin (17 mice, 155 sessions). To ascribe excess effects to trials with different outcomes, we pooled trials either within 0.5s bins of time with respect to the cue on trial n (as in **Fig 1g**). We also tried more broadly pooling early and rewarded trials (statistics reported in-line in the text). “Early trials” included all trials with a first-lick time falling within a “notch” of the timing distribution (between the local median and opening of the reward window at 3.333s). We looked at notch trials because we expected learning updates (late-shifting) and R2M effects (early-shifting) for these trials to be in opposite directions, presenting the most challenging conditions to detect learning effects. To put pooled early and rewarded trial analyses on the same footing, the rewarded trial “notch” included trials pooled within an equivalent interval in the reward window (e.g., if the notch were between 3 to 3.333s for a local measurement, the rewarded notch interval was taken as 3.333-3.666s).

All three R2M approximation methods (A, B and C) returned similar results in all analyses. R2M effects were present in the dataset, indicating the mice did indeed have a tendency to “regress to the median” movement time, consistent with a stochastic generative process underlying timing. However, ΔT was in excess of R2M by all metrics and depended on the trial outcome: early trials (>0.5 to 3.333s) showed late-shifting in excess of chance, whereas rewarded trials showed early-shifting—consistent with timing updates in excess of chance. Interestingly, “reaction” trials, within 500ms of the cue, also showed small but significant early-shifting, which was likely a multi-trial effect (i.e., rewarded trials were often followed by reactions, and the subsequent trial (n+2) also appeared to show some residual early-shift). Although there was no opportunity for mice to be “late” in this version of the timing task, rare trials in which the first-lick fell within the ITI (unrewarded) showed subsequent late-shifting, consistent with DA dips observed on these trials. It is important to note that these late-shifts represent shifts in the *median* of the putative latent distribution on trial n+1, which were small (100s of milliseconds) compared to the first-lick time (>7s)—which means that the first-lick time on trial n+1 was still more likely to be much earlier than that on trial n when trial n’s first-lick occurred during the ITI. Finally, we note that the size of the excess ΔT effects was on the same order of magnitude as one-shot optogenetic effects (tens to hundreds of milliseconds, **Fig 3**), suggesting that outcome-related DA updates the stochastic “set-point” of the latent timing distribution on trial n+1.

### Photometry

#### Signal collection

Fiber optics were illuminated with 475nm blue LED light (Plexon, TX or Doric Lenses Integrated Photometry Cube, Quebec, Canada) (SNc/VTA: 50μW, DLS: 35μW, VLS: 30µW) measured at patch cable tip with a light-power meter (Thorlabs, NJ). Green fluorescence was collected via a custom dichroic mirror (Doric Lenses) and detected with a Newport 1401 Photodiode (Newport Corporation, CA) or integrated Doric cube. For red florescence, 550nm lime LEDs were used (Plexon or Doric Lenses Integrated Photometry Cube). Fluorescence was allowed to recover ≥1d between recording sessions. To avoid optical crosstalk, the optical control channel (e.g., tdt or GFP) was recorded at one of the other implanted sites while GCaMP6f, dLight1.1/1.3/rdLight1 or DA_2m_ were recorded simultaneously only at the other implanted sites.

**dF/F.** Raw fluorescence for each session was pre-processed by removing rare singularities (single points >15 STD from the mean) by interpolation to obtain F(t). To correct photometry signals for bleaching, dF/F was calculated as:

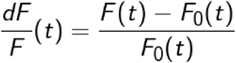

where F_0_(t) is the mean fluorescence during the 5s preceding the current trial, as well as 5 trials before the current trial and 5 trials after the current trial (as we characterized and validated previously^7^).

#### Correlation of baseline photometry signals with first-lick time

Baseline signals were de-noised by pooling based on first-lick time into 0.25s pools from 0.75 to 4s after the cue, as in **Fig 1d**. Pools with first-lick times after 4s were not included due to insufficient numbers of trials in each pool to de-noise the measurement (inclusion criteria >=175 trials). Pearson’s r was calculated from each pool’s average dF/F signal calculated within windows of time relative to lamp-off (left of **Fig 1d**, r: –0.63; –10.5 to 10s before lamp-off) or the cue (right of **Fig 1d**, r: –0.79; –0.4 to 0s before the cue (excludes lamp-off event)). These signals were related to the earliest first-lick time in each pool.

#### Generalized Linear Model (GLM)

To test the independent contribution of each task-related input to the photometry signal and select the best model, we employed nested fitting, in which each dataset was fit multiple times (in “nests”), with models becoming progressively more complex in subsequent nests. The nests fit to the photometry data employed the inputs X^(j)^ at each *j*^th^ nest:

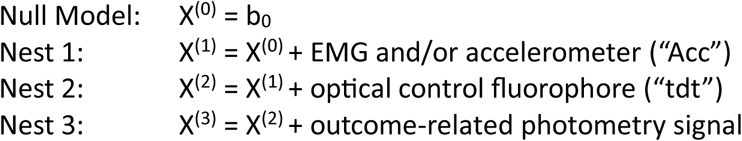

The outcome-related photometry signal was taken as the mean dF/F signal within the first 500ms after the first-lick (peak or median dF/F produced similar results). All *d_j_* design matrix predictors were 0,1 normalized.

The GLM for each nest took the form:

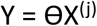

Where Y is the *1*x*N* vector of the change in the movement time between the current trial, n, and the next trial, n+1 (ΔT_(n+1)-n_), where *N* is the total number of trials in the session; X^(j)^ is the *d*x*N* design matrix for nest *j*, where the rows correspond to the *d_j_* predictors for nest *j* and the columns correspond to each of the *N* trials of Y; and ϴ is the *d*x*1* vector of fit weights. Because of day-to-day/mouse-to-mouse variation (ascribable to many possible sources, *e.g.*, different neural subpopulations, expression levels, behavioral states, *etc*.), each session was fit separately.

To guard against a trivial relationship between the trial outcome and DA signal, separate models were fit for early or rewarded trials.

Among the nests, the best model for each session was selected by BIC (AIC and AICc were typically in agreement with BIC), which was typically the model including the photometry signal. All reported data is for the full model.

#### Residual GLM (Extended Data Fig 4a)

To determine whether outcome-related DA was predictive of ΔT in excess of chance (regression to the median effects), the next trial’s movement time was shuffled, and the GLM was refit on the shuffled data (10,000 shuffled datasets). The coefficients from the 50th percentile shuffled model (50th percentile shuffled DA coefficient) were used to predict the true data (y-fit shuffle) and calculate shuffled-R^2^. Y-fit shuffle was subtracted from the true dataset to obtain a residual dataset, and the residual model was derived by refitting the GLM on the residual dataset.

#### Shuffled models/regression to the median (Fig 2e, Extended Data Fig 4b-c)

To guard against spurious relationships between DA and ΔT arising from behavioral non-stationarity, each session was divided into four quartiles of trials (same number of trials with participation in each quartile) to isolate more stationary periods of behavior. The GLM was fit on each quartile (including movement/optical artifact control signals and mean DA photometry signal during the first 500ms after the first-lick). The trial order (for each quartile) was then shuffled 10,000x and the GLM was fit on each shuffled dataset to produce the distribution of DA coefficients for each quartile assuming no trial-order dependence. To determine if the true fit coefficient was in excess of the shuffle, a Wilcoxon rank sum test was applied to compare the set of true coefficients across quartiles (and sessions) to the set of mean shuffled DA coefficients.

#### Standard box plots

Throughout the paper, whenever used, boxplot centerline: median; box limits: upper and lower quartiles; whiskers: 1.5x interquartile range; points: individual datapoints, including outliers.

### Bleaching analyses

As is standard for photometry measurements, we corrected for bleaching using dF/F. Nevertheless, there remains some possibility that bleaching is either not fully corrected by this processing *or* that dF/F itself introduces some insidious artifact that could explain the downtrend in rewarded DA transients (“rewDA”) across the session. *A priori*, this is unlikely:

1. The amplitude of unrewarded transients showed relatively little change across the session (**Extended Data Fig 7d,j**)—bleaching artifacts would likely affect signals similarly for all trial outcomes.
2. Indirect pathway striatal projection neuron (iSPN) GCaMP signals showed the *opposite trend* across the session on average: rewarded iSPN transient amplitude *increased*, on average (n=3 A2A-cre mice, 30 sessions; **Extended Data Fig 8 bottom row**). iSPNs receive negative input from DANs, thus these findings are consistent with decreased DAN signaling.
3. The timing set-point showed abrupt shifts that occurred at the moments of the greatest decline in rewDA (**Fig 4e**), these changes occurred at different times in the session (**Extended Data Fig 7n**) and occurred significantly earlier in the No Juice Task (**Fig 4k**). Bleaching would likely cause similar decay rates across all sessions from the same mouse.
4. Reward-related tdTomato transients (rew-tdt) did not show the same pattern of decline across the session (**Extended Data Fig 7f,k, Extended Data Fig 8 top row**) and did not explain abrupt shifts in set-point (**Extended Data Fig 7h**).
5. Causal effects of rewDA on set-point cannot be explained away by bleaching (**Fig 5, Extended Data Fig 9**).
6. Rapid Satiation (**Fig 5g-i**) caused a sudden, precipitous decline in rewDA that was significantly different from that observed in control sessions in the same mice, which cannot be explained by bleaching.

Nevertheless, to directly assess the contribution of bleaching to the decline in rewDA, we derived a GLM predicting the peak amplitude of rewDA from bleaching-related and behavioral predictors. To derive a bleaching predictor, we applied the raw fluorescence, F0, from the dF/F correction. F0 was measured for each trial as the average fluorescence signal during the 5s preceding the Lamp-off event, averaged across the 10 nearest neighboring trials.^7^

We first fit the model with F0 alone. As expected, because F0 decreases monotonically, F0 was positively correlated with set-point and the decline in rewDA, and thus was a significant positive coefficient in the “bleaching-only model.” This verified that bleaching was encoded properly.

Subsequently, we introduced behavioral predictors to derive nested models containing permutations of the following predictors:

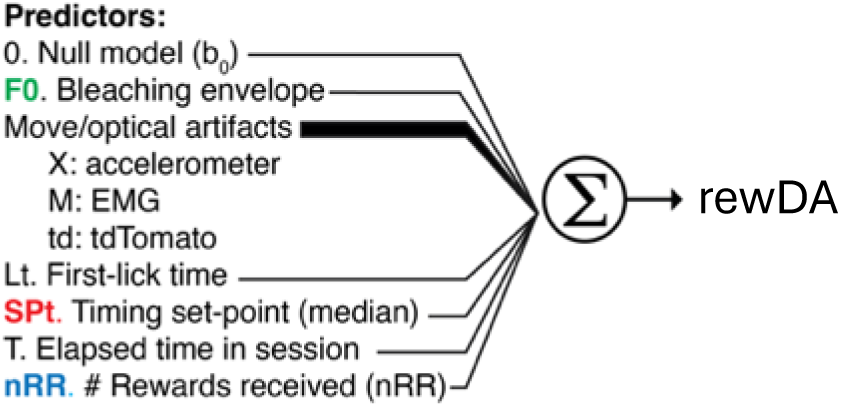

Across DAN/DA recordings, predictors related to ongoing body movements, optical artifacts, and first-lick time were relatively poor predictors of rewDA, as evaluated by variance explained and 4 complementary model criterion metrics (AIC, BIC, AICc, and CAIC). For the +Juice Task, the best models always included the number of rewards received (nRR) as a significant predictor, which weakened F0’s coefficient, indicating that variance in rewDA was independently explained by variables related to satiation and/or behavioral accuracy, in excess of bleaching. Supporting this, when the models were fit on data from the Timeshift Task, in which the set-point no longer trivially correlated with nRR, set-point became a stronger predictor in the model, and all 4 information criterion metrics agreed that the most parsimonious model of rewDA required both set-point and nRR as significant predictors, in excess of bleaching. Together, these data indicate that rewDA signals contained information related to propensity to move +/-satiation, in excess of bleaching. Thus, bleaching did not explain away the relationship between rewDA and changes in the timing set-point.

### Optogenetics

#### Opsin expression

3 cohorts of mice were tested: Activation (ChR2, 4 mice), Inhibition (stGtACR2, 4 mice or JAWS, 1 mouse), and No Opsin (no SNc injection, 4 mice). All mice co-expressed rdLight1 or GFP at VLS for co-monitoring of stimulation effects on DA release. All stimulations were delivered via fiber optic cannulae implanted over SNc.

#### Co-monitored photometry

We selected ChR2 (blue-activated opsin)/rdLight1 (green-excited fluorophore) as our DA co-monitoring strategy because we had previously observed issues with off-target excitation of red light activated opsins (e.g., ChrimsonR) during prolonged co-recording of dLight1.1 (a blue light-excited fluorophore) in DLS (red opsins are also activated by blue light, albeit with lower efficiency, which becomes significant during hours-long timecourses of photometry illumination in our behavioral sessions).^7^ This off-target excitation would have created a confound, because we had previously shown that DAN activation limited to the self-timing interval increases the probability of early first-licks.^7^ ChR2, by contrast, is not appreciably activated by green light (used for rdLight1 excitation), and was thus a cleaner preparation. One challenge of blue stimulation/green-excited photometry is that the stimulation light causes a DC-offset artifact in the red photometry channel while the stimulation LED is turned on. To avoid crosstalk between the stimulation LED and the photometry recording, two complimentary correction methods were used, with similar results: 1) the brief stimulation up-times were omitted from the photometry signal (missing points filled by interpolation), or 2) the signal was decontaminated by measuring the instantaneous increase in rdLight1 signal at the beginning of each pulse of blue light, which was subtracted from the voltage trace during each pulse. Both corrections were taken before dF/F correction and yielded similar photometry signals, however the second method had the advantage of compatibility with continuous illuminations used during stGtACR2 inhibition experiments (see **Stimulation Parameter Calibration**, below).

#### Stimulation parameter calibration

Stimulation parameters were initially calibrated outside the context of the task. Mice sat on the typical behavioral platform and received 10 unexpected juice rewards (ca. 4µL each), spaced at least 15s apart. A grid screen of stimulation parameters and light powers identified conditions that best recapitulated both the amplitude and width of the typical reward-related transient. Mice received 10 stimulations with each stimulation protocol, and DA release in VLS was monitored throughout the calibration procedure with rdLight1. Light intensity and pulse width were varied until the resulting rdLight transient matched the amplitude and width typically seen with unexpected juice rewards.

The final parameters used in activation experiments were as follows: Optogenetic stimulation was delivered bilaterally via blunt fiber optics at SNc (5mW 465nm LED, 10Hz, 250ms, 35% duty cycle). One ChR2 animal had partial response to 5mW blue light and thus received 10mW blue light, with a larger but still partial effect (Mouse X4-3-1M). One mouse from the pilot study (X4-1-1M) used different parameters for its first four sessions (10Hz, 500ms, 20% duty cycle), which produced slightly wider DA transients but behavioral effects consistent with the optimized parameters.

We also attempted to inhibit DANs (n=4 stGtACR2 or n=1 JAWS), but despite serious effort (5 mice), we could not identify stimulation parameters that would prevent reward-related DA transients, despite finding parameters that caused DA dips outside the context of the task. Furthermore, more aggressive inhibition at higher light powers (e.g., 5-30 mW) in the context of the task—but not outside the context of the task—caused *seconds-long* rebounds in baseline DA after the offset of inhibition (i.e., this was not the typical transient “rebound spike” sometimes seen after inhibition, but a more sustained increase in DA). Thus—paradoxically—aggressive inhibition tended to *raise* DA overall on stimulated trials. We suspect inhibition was ill-posed in this preparation: we could not initiate stimulation fast enough after detecting the lick to prevent reward-related action potentials from triggering DA release, and more powerful inhibition caused paradoxical rebounds. Others have also been unable to effectively block reward-triggered DA transients with inhibition and have proposed similar rationales (e.g., Joshua Dudman, personal communication, and Ref ^32^).

Because we could not prevent first-lick related transients, we did not proceed with the Inhibition cohort. Instead, in lieu of optogenetic inhibition experiments, we caused reductions to outcome-related DA through non-optogenetic manipulations (**Fig 4f-l, Fig 5**), whose effects were consistent with the GLM.

#### One-shot stimulation analyses

Initial power simulations indicated that ∼5,000 trials (10x 500 trial sessions) would be necessary to achieve 80% power to detect a 20% change in first-lick time. Thus, at least 10 sessions of stimulation (∼500 trials each) were collected from each animal.

Three complementary analyses examined single-trial stimulation effects (**Extended Data Fig 6**):

**1. “deldel,”** difference in the timing update (del1) between stimulated vs unstimulated trials (del2): Δ(ΔT(n+1)–n)stim-unstim=ΔT(n+1)–n | trial n stimulated - ΔT(n+1)–n | trial n unstimulated
**2. “dAUC,”** difference in the area under the first-lick empirical continuous distribution function (eCDF) curve of the next trial (n+1) timing distribution for stimulated vs unstimulated trials (as described previously^7^).
**3. “cdf-rank,**” percentage of points in the n+1th trial eCDF after stimulated trials that were less than matching points in the n+1th trial eCDF after unstimulated trials. eCDFs were interpolated such that each point was matched between the stimulated and unstimulated curves.

Analyses were performed at 3 levels, with consistent results across analyses and levels: <u>By Session</u>

*Within single session analyses*: deldel, dAUC and cdf-rank analyses compared the distributions or first-lick times on the next trial after stimulated vs unstimulated trials. As expected from preliminary power calculations, these statistics were noisy (**Extended Data Fig 6 top row, single dots**).

*Between groups single session comparisons:* The test statistic for each single session was bootstrapped 10,000x, and the mean of the single session test statistics for each group were subtracted to derive a between groups comparison test statistic (activation – no opsin), which was then compared by Wilcoxon rank sum test (**Extended Data Fig 6 top row, between group statistic**).

#### By Mouse

*Within “Mega” session comparisons*: Because initial power simulations suggested 5000 trials were needed for 80% power to detect a 20% change in first-lick time, data were pooled across stimulated sessions from individual animals to generate “mega” sessions, which had ∼5000 trials each. The test statistic for each mega session was reported as dots in **Extended Data Fig 6, second row**.

*Between groups mega session comparisons*: The test statistics for each group of animals’ mega sessions were compared by Wilcoxon rank sum test. 2-sided unpaired t-test found similar results.

#### All Mice

*Within “super mega session” comparisons*: All data from all sessions in each cohort were pooled to generate a “super mega session” to compare all stimulated and unstimulated trials by 10,000x bootstrap procedure (same as used on within-single session analyses). Test statistic and p-value reported in the Main Text and **Extended Data Fig 6, third row**.

There were 3 instances where sessions had to be excluded from some analyses: 1) Due to a software error, two sessions (Lazarus_VLS_25 stim, Lazarus_VLS_34 sham) recorded too few trials to compute single-session cdf-based bootstrapped comparisons (dAUC and cdf-rank) and were omitted from those analyses. The error was not detected until after the animal had retired. 2) Partial stimulation effect: 5mW light power was insufficient to elicit DA release in one animal (X4-3-1M). The light power was subsequently increased to 10mW, which produced an improved partial effect on DA release. Only 10mW sessions from this animal were used for analyses. 3) One animal (X4-3-2M) suffered from a dermatological condition that developed over the course of the experiment, which reduced continuity with the grounding plate, resulting in inconsistent lick detection on a subset of sessions—out of caution, affected sessions (defined as having >5% of trials with failed lick detection) were excluded. After veterinarian treatment and recovery, the animal was implanted with a subcutaneous ground wire, which corrected lick detection. Sessions recorded with the ground wire were included in the analysis.

### Whole session behavioral analyses

#### “Set point” calculations: Moving median first-lick time

For visualization, the timing set-point was defined as the 50-trial moving median centered at the current trial, excluding missing trials (e.g., no-lick) and reactions (first-lick <0.5s). This captured the peak of the timing distribution (whereas including reactions tended to underestimate the peak). For quantitative comparison to rewarded DA transients (which occurred only on a subset of trials) and in tasks with rapidly changing set-point dynamics, the number of trials included in the moving median was adaptively reduced to avoid over-smoothing (10-50 trials used, depending on analysis: e.g., Timeshift No Juice task used 15 trials). Data provenance text files are included with each figure and specify the smoothing used in each analysis.

#### +Juice/No Juice Task Comparisons

For animals with No Juice Task sessions (n=5 VLS rdLight mice, 11 sessions), the nearest +Juice session with photometry was matched for comparisons of behavior (timing set-point) and outcome-related DA signals. For set-point analysis, 15 sessions were available for comparison (e.g., **Figs 1b, 4b-d**). Some flanking sessions were collected without photometry—in these situations, the most recent photometry session was used for DA analyses. Together, this produced 18 paired photometry sessions for No Juice vs +Juice comparisons (**Fig 4i-k**; n=5 mice). Except for one session (Randy_VLS_47), paired +Juice photometry sessions happened before the paired No Juice session. Because several +Juice sessions only recorded 500 trials total, for consistency, we used only the first 500 trials from each session for paired comparisons. The Q1 and Q4 lick-triggered averages shown in **Fig 4** were thus taken on trials 1-125 and 376-500, respectively, irrespective of session duration.

One No Juice session (Paisley_VLS_23) was not usable for photometry analysis due to power loss of the photodetector midway through the session, and one paired +Juice photometry session was excluded due to issues with the photometry light path during data collection (Elle_VLS_35); these sessions were still used for behavioral analyses. Paired +Juice sessions (15 total) were taken from the set of flanking +Juice sessions. These +Juice sessions showed no consistent differences in set-point dynamics (indeed the variance in +Juice set-point dynamics was lower than that observed in the No Juice Task). Nevertheless, we note that two of these +Juice sessions were SNc stGtACR2 sessions that showed no effect of stimulation on co-monitored DA release. Out of caution, trials with stimulation were excluded from DA analysis, though, importantly, inclusion of these sessions was conservative because inhibition was expected to cause late-shifting, which would have obscured differences between the No Juice and +Juice Tasks. Furthermore, paired No Juice Task sessions still had significantly lower rewarded DA transient amplitudes and were significantly late-shifted compared to these two sessions. Of the remaining paired +Juice sessions, seven contained rewarded trials that delivered different volumes of juice: either the usual amount (∼4µL, 60% of trials), low juice (∼1µL, 20% of trials) or high juice (∼13µL, 20% of trials), where juice level was randomly interleaved among rewarded trials. Rewarded DA transient amplitudes were similar for all juice levels, on average, but the width of transients was systematically different (higher volume/wider transient). Thus, out of caution, rewarded trials with either low or high juice in these sessions were excluded from DA analyses.

#### Set-point cross-over analysis (Fig 4e)

Sudden shifts of the set-point into the reward window were eventually observed in all animals when allowed to work ad libitum, typically between trials 400-1100. However, because many sessions were only recorded for 500 trials, only a subset of sessions were recorded long enough to observe cross-overs (we validated this in a large behavioral dataset from animals allowed to work until disengagement, 700-1800 trials, 3 mice, 120 sessions). Sessions showing set-point cross-over were identified in the SNc GCaMP or VLS rdlight1 datasets as those sessions with set-point non-stationarity of >=+1.5s (16 sessions total). If the set-point started within the reward window at the beginning of the session, the set-point had to first exit and then re-enter the reward window to be considered a cross-over event (1/16 sessions). Evolution in first-lick time and rewarded DA transient amplitude (“rewDA”) was taken as a moving median over 75 trials (37-38 either side of each trial), where missing data was interpolated linearly (edges excluded as above to avoid artifacts). To de-noise the derivative of the rewDA signal, rewDA was first downsampled (every 15 trials taken). The downsampled trial with maximum derivative of the DA signal (–1*rewDA’) was compared to the “set-point cross-over trial,” defined as the trial on which the median of the timing distribution first crossed into the reward window (>3.333s).

### Cued Reaching Task

#### Behavior and analysis

dLight1.1 fiber photometry recordings were taken from pDMSt during the cued reaching task as previously described.^63^ zdF/F was calculated over a 30s running baseline window. Reaches were determined from the video, as described in Ref^63^. The metric of DA for rewarded trials was encoded as the single-trial maximum signal within a 12s window starting 1.5s before the reach, minus the average baseline signal over the 1.5s interval preceding the reach. Because miss trials, defined as a reach when the pellet was omitted, were not associated with transient DA increases, the metric of DA on miss trials was designated to capture sustained DA signals. The metric of DA on miss trials was the mean signal 0.95 to 2.7s after the cued reach (these means were not baseline subtracted). ΔT-reach was defined the reaction time on trial n+1 minus the reaction time on trial n. A minimum ITI of 18s was enforced to ensure a several second delay between cessation of food pellet chewing and the next trial-start. Ongoing body movements/optical artifacts were controlled for with a movement predictor, computed as a robust z-score of the absolute derivative of video pixel intensity within the video zone overlapping the position of the paws before the start of a reach. GLMs of the form ΔT-reach ∼ DA + movement + b_0_ were fit on 0-1 normalized predictors. Single session models were fit on all sessions with >20 trials of each category of trial. Across session-pooled trial regressions were fit on 0-1 normalized predictors normalized within each session.

### Code Availability

All custom behavioral software and analysis tools are available with sample datasets at https://github.com/harvardschoolofmouse.

### Data Availability

This study used both publicly available datasets that we previously released (Zenodo; DOI: 10.5281/zenodo.4062748), as well as original datasets. All original datasets supporting the findings of this study will be made publicly available at the time of publication via DANDI, with source data files provided for all figures.

## Acknowledgements

We thank L. Hou, E. Sayed, J.G. Mikhael, and S. Hrvatin for discussions; M. Yang, S.L. Jacobs-Skolik, N. Hunter, V. Berezovskii, J. LeBlanc, T. LaFratta, O. Mazor, and P. Gorelik for technical assistance; and C. Harvey, S.R. Datta, M. Andermann, A. Mohebi and W. Regehr for reagents. The work was supported by NIH grants UF-NS108177 (J.A.A), U19 NS113201 (J.A.A, B.L.S) and K99 MH127471

(K.R.); NIH core grant EY-12196; the Valhalla Foundation (A.E.H.); the Whitehead Institute Innovation Initiative (A.E.H.); the Whitehead Fellows Program (A.E.H.); and the Helen Hay Whitney Foundation postdoctoral fellowship (K.R.). The funders had no role in study design, data collection and interpretation, or the decision to submit the work for publication.

## Author contributions

**A.E.H. and J.A.A.** conceived the study, designed the experiments, acquired funding, supervised the project, and wrote and edited the manuscript. **A.E.H.** Performed experiments, wrote the code for analysis, analyzed the data, and wrote the first draft. **I.C.W., Q.D., Z.O. and P.A.** assisted with experimental design, performed experiments, processed data, and assisted with analysis. **E.N.** performed experiments. **K.R. and B.L.S.** contributed data and analyses from the cued reach task. All authors reviewed the final manuscript.

## Additional Information

Corresponding Author: Allison E. Hamilos

## Supplementary information

Extended Data Figures

---

**Extended Data Fig 1.**
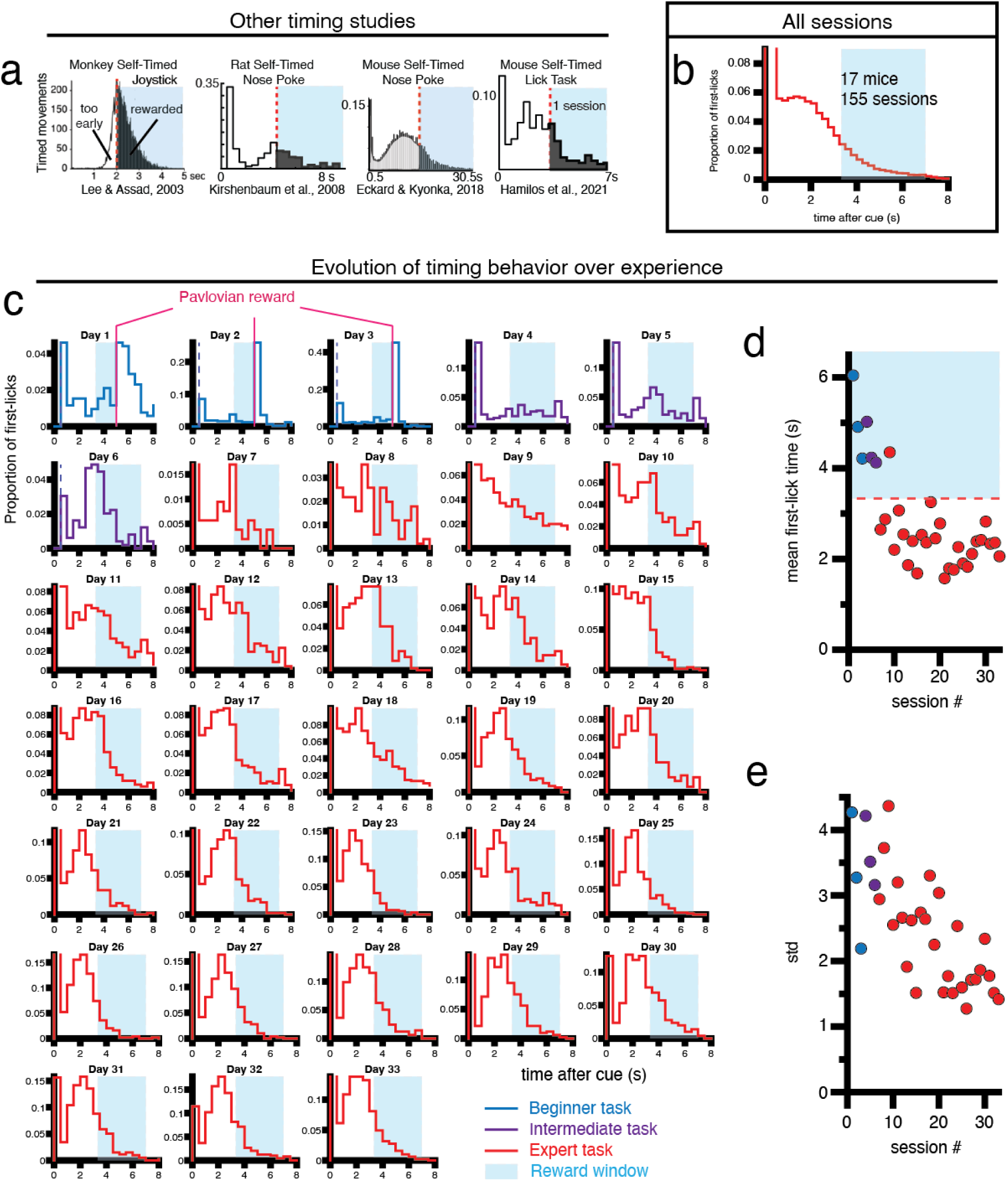
Timing behavior. a, Timing distributions from monkeys, rats, and mice from other studies show early-bias relative to the reward window. Right-most panel is from our previous study using the self-timed movement task. Figure panels/data adapted and used with permission. **b,** Timing distributions across all mice and photometry sessions (including day 5 of training onward), all trials in each session included. **c,** Histograms show the distribution of first-licks after the cue on each day of training with the self-timed movement task (one representative mouse; additional examples of training progress shown in Hamilos et al., 2021 Figure 1—figure supplement 1). This mouse was trained for 33 sessions with 500 trials/session, with no training days on weekends. All mice underwent three stages of training (“Beginner,” “Intermediate” and “Expert” levels), each of which is described in detail in the Methods. Briefly, the Beginner Task encouraged participation and learning in two ways: 1) with a permissive 0.5s reaction window after the cue, such that if the mouse licked immediately after the cue, it was allowed to respond again on the same trial, and 2) an automatic “Pavlovian” juice reward at 5s *if* the mouse had not already licked between 0.5-3.3s after the cue. The Intermediate Task was the same as the Beginner Task, but there were no longer automatic Pavlovian rewards delivered at 5s. The Expert version was the “full” version of the task, in which there was no longer a permissive reaction window. All data in the paper (except for this panel) are shown from animals performing the Expert version of the task. The mouse shown here was transitioned from the Beginner Task (blue traces) to the Intermediate Task (purple traces) on day 4 of training. Transition to the Expert Task (red traces) occurred on day 7. Y-range of histograms was maximized to clearly show the timing distribution peak; an initial reaction peak (<0.5s) was truncated in some plots and indicated by open trace. First-lick timing distributions became stable by day 20 of training, and early timing bias persisted throughout training for all animals when juice rewards were used. **d,** Mean first-lick time for each day of training. Early timing bias (mean first-lick time between 2-3s) is firmly established by day 10 and persists throughout training. **e,** Although the mode of these distributions remained early-shifted throughout training, the standard deviation of first-lick time decreased as the timing distribution became more stereotyped over the course of training.

**Extended Data Fig 2.**
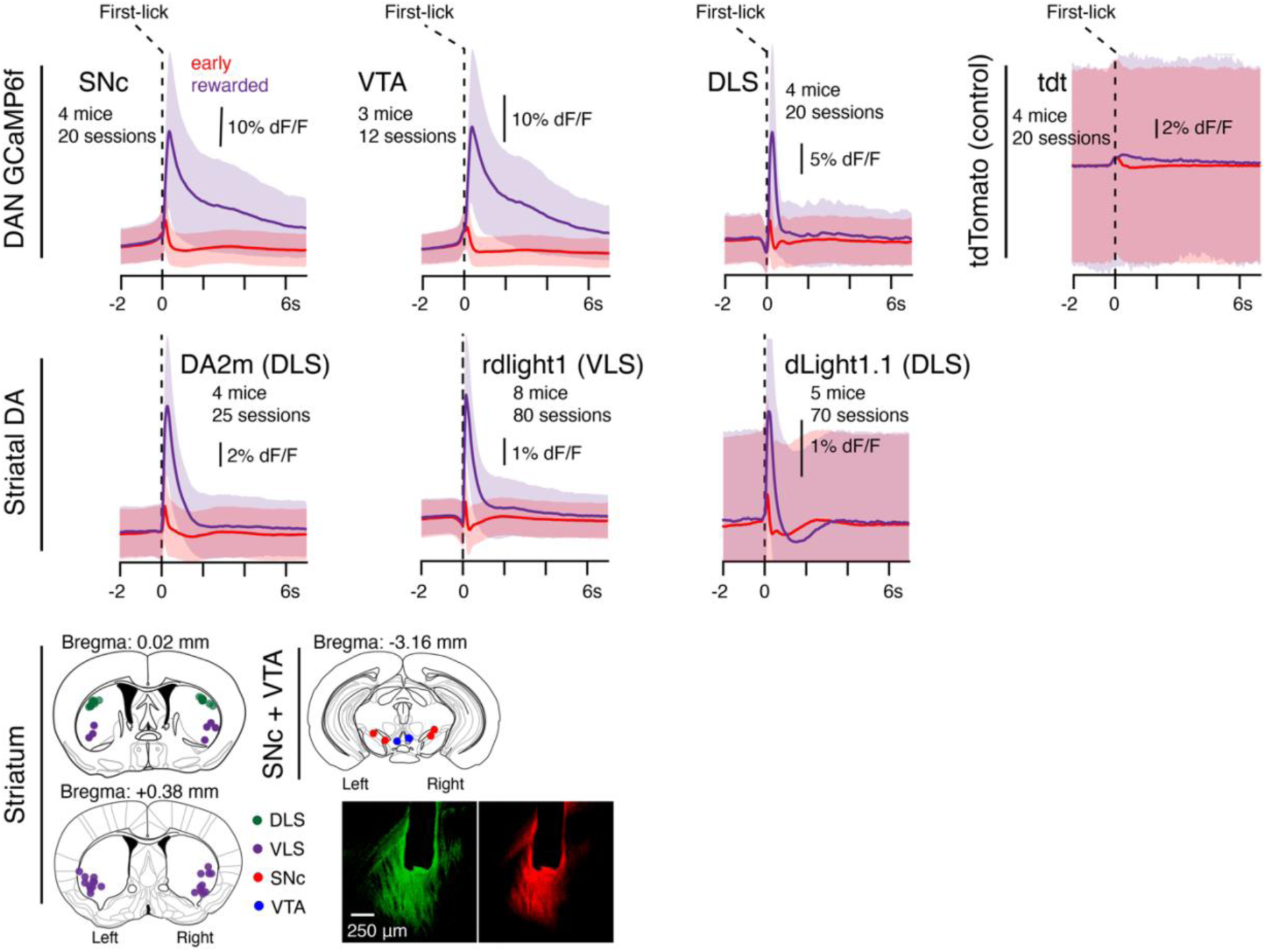
Mean outcome related dopaminergic photometry signals. Shading: standard deviation. Inset: Recording locations for dopaminergic fiber photometry recordings and example epifluorescence from a coronal brain section of a dLight1.1 cohort mouse, which shows the blunt fiber tip and co-expression of the DA indicator and control red fluorophore in the DLS.

**Extended Data Fig 3.**
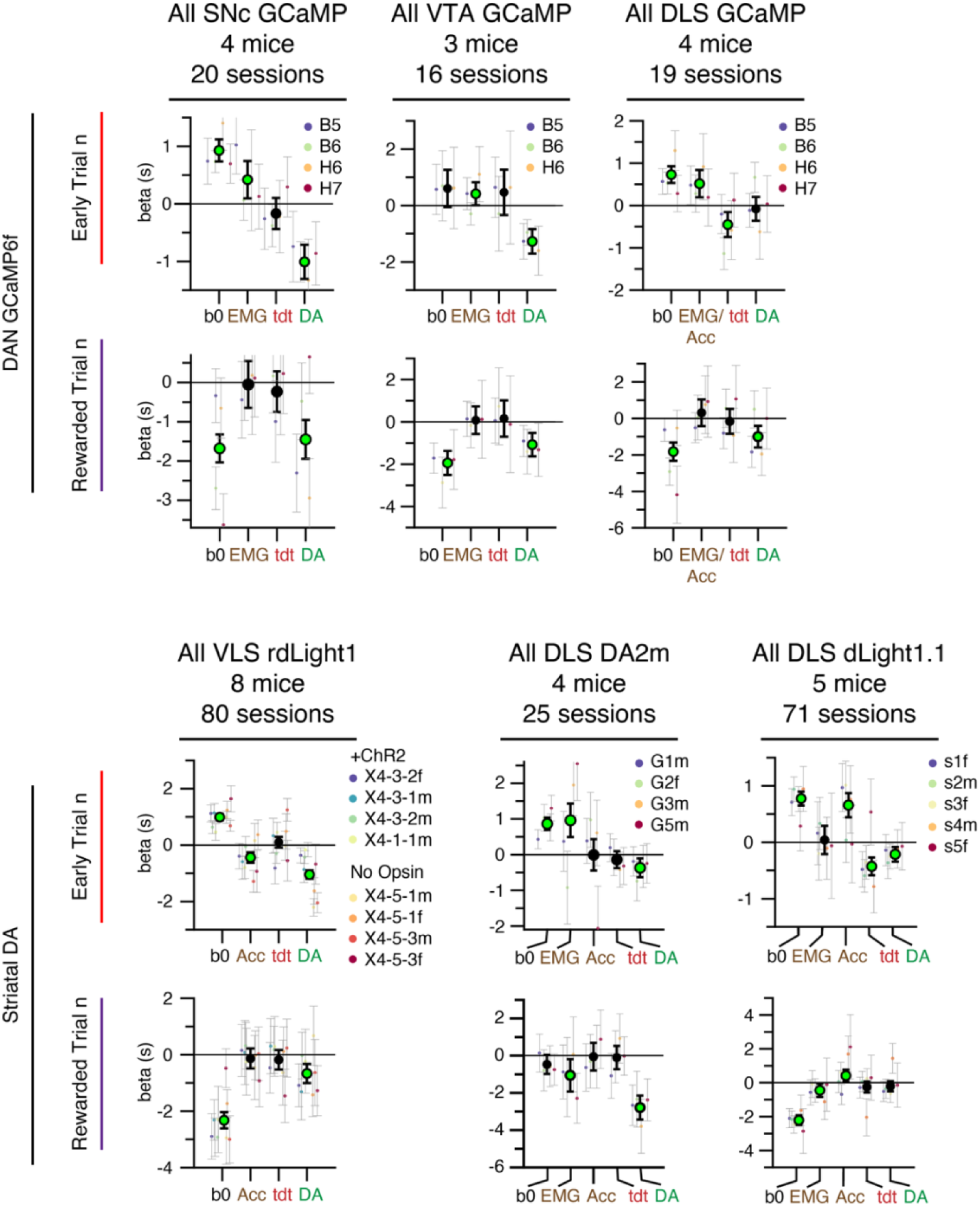
GLM coefficients, all signals, all sessions, all mice. Small colored dots indicate coefficients from individual mice. “Acc:” accelerometer signal (present in some cohorts of animals), “EMG/Acc:” sessions had either EMG or accelerometer present; the signal that was present was used for this predictor. Note that the optical control channel “tdt” in dLight1.1 recordings was RFPturbo, and one rdLight1 mouse (x4-1-1m) used mCherry instead of tdt.

**Extended Data Fig 4.**
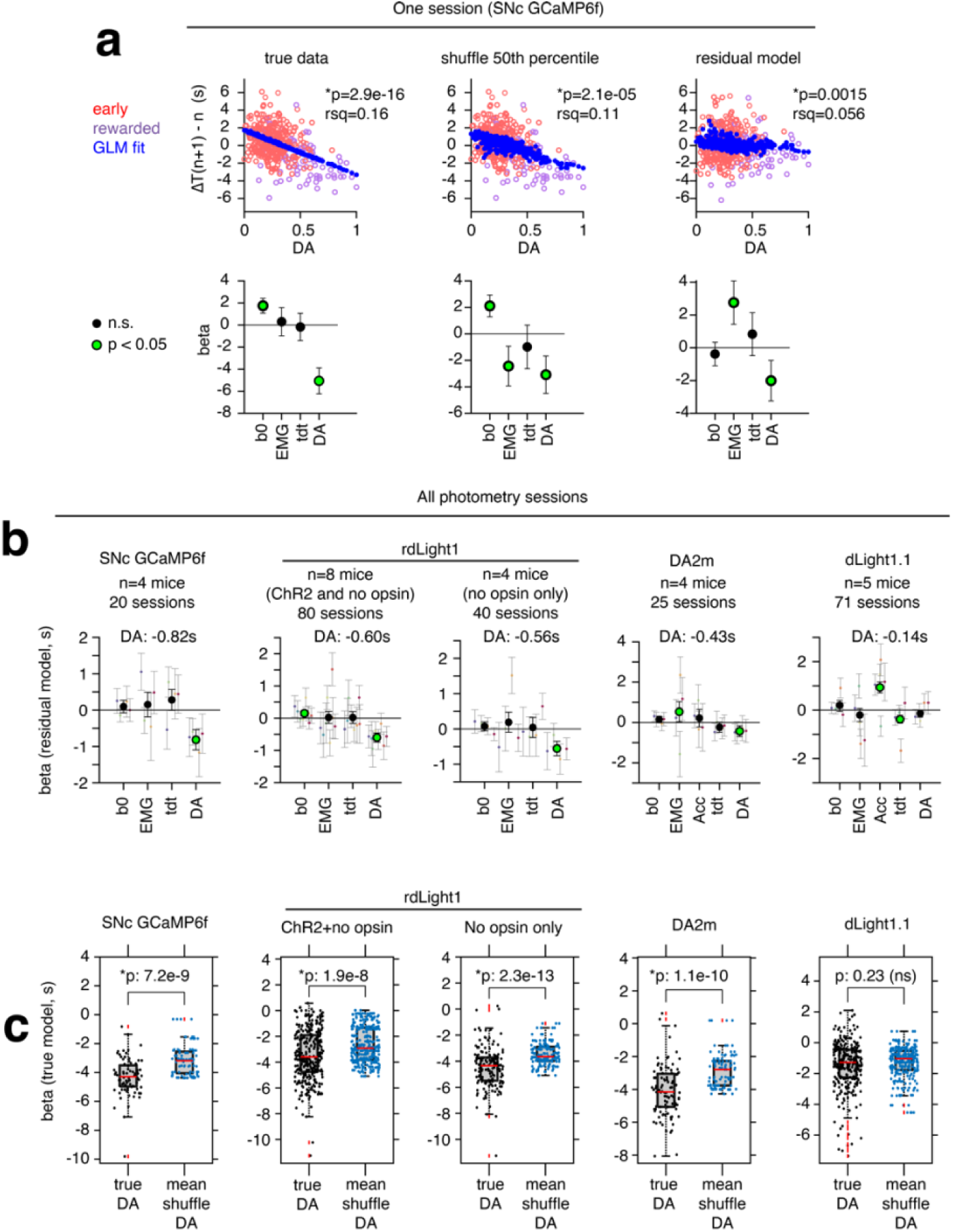
Outcome-related DA explains ΔT in excess of chance. a-b, Shuffle models. The next trial’s movement time was shuffled, and the GLM (**Fig 2a**) was refit on the shuffled data (10,000 shuffled datasets total). The coefficients from the 50^th^ percentile shuffled model (50^th^ percentile shuffled DA coefficient) were used to predict the true data (y-fit shuffle) and calculate R^2^ (panel *a*, middle). Y-fit shuffle was subtracted from the true dataset to obtain a residual dataset, and the residual model was derived by refitting the GLM on the residual dataset (panel *a*, right). Outcome-related DA remained a significant predictor to the residual model, both in single sessions (panel *a*, right-bottom panel) and across sessions for SNc GCaMP, VLS rdlight1, and DLS DA2m. DLS dlight1.1 DA signals did not remain significant after the shuffle procedure, likely due to relatively poor S:N quality in the dlight1.1 dataset. **c,** Comparison of true vs shuffled model DA coefficients fit on quartiles of trials in the session. To guard against spuriously significant DA coefficients arising from non-stationarity in timing behavior across the session, true and shuffle models were fit on quartiles of trials in each session (e.g., first quarter of trials in the session, second quarter, etc.). Consistent with the residual models in panel *b*, the population of true model DA coefficients was significantly larger in magnitude (more negative) than the population of mean shuffled model DA coefficients (except for DLS dlight1.1; Wilcoxon rank sum test, *p<0.05). Outcome-related DA is thus predictive of ΔT in excess of regression to the median effects (see Methods) and unlikely to be explained by behavioral non-stationarity.

**Extended Data Fig 5.**
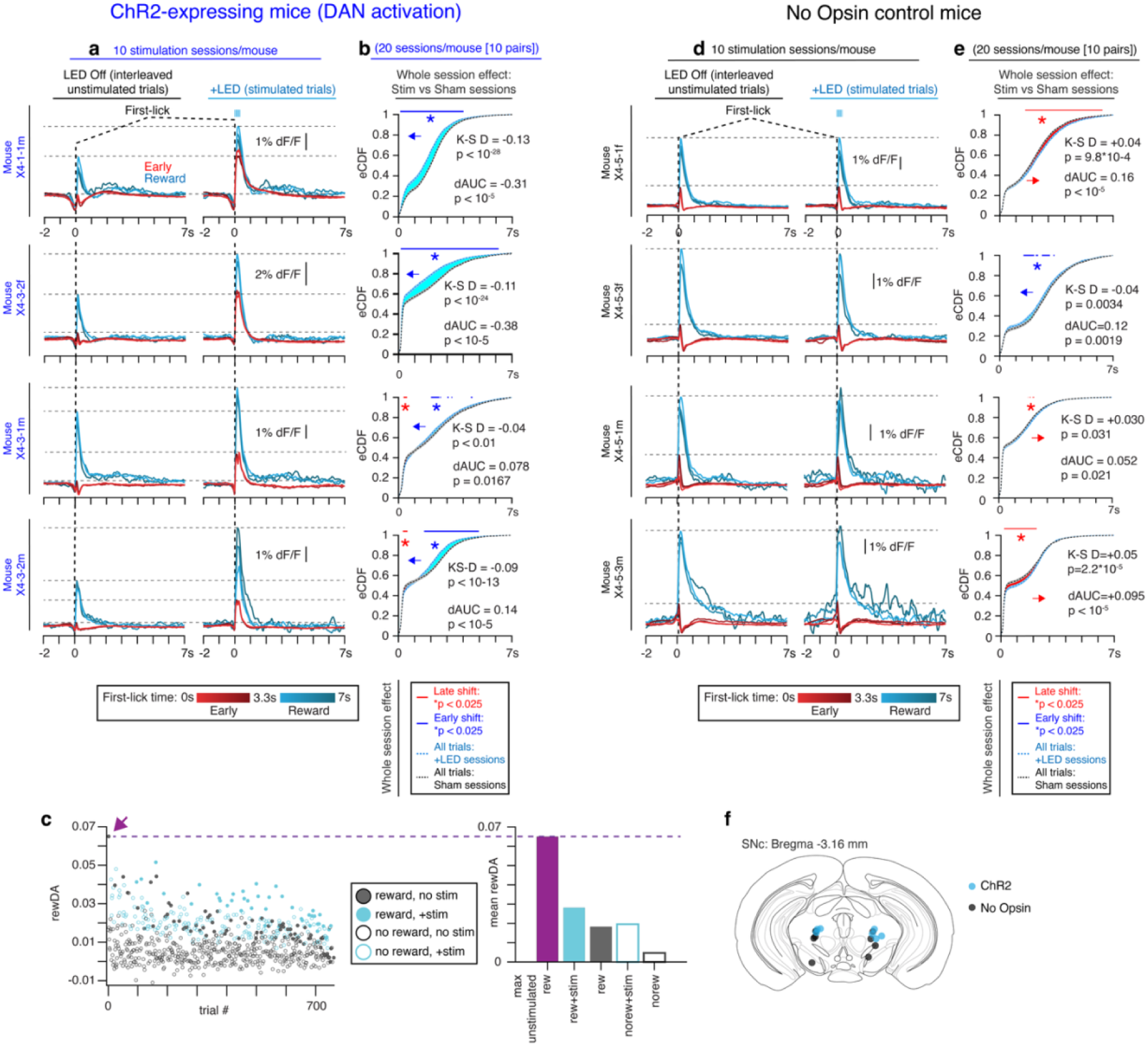
One-shot optogenetic manipulations—all mice. a, VLS rdlight1, all ChR2 stimulation sessions, 10 sessions/mouse. Each row is a different mouse. LED was triggered on 30% of randomly interleaved first-licks. **b,** Comparison of trial n+1 first-lick distributions (all trials) from stimulation sessions (blue trace) vs interleaved sham sessions (black trace). Blue shading: early-shift effect of LED; red shading: late-shift effect. Arrow: indicates overall direction of effect. Top bars (blue/red): denotes significant difference between eCDF curves (*p<0.025, 95% CI, 1000x bootstrap test). “K-S D”: signed Kolmolgorov-Smirnov distance metric (negative indicates early shift effect). “dAUC”: difference in area under eCDF curves, p: 10,000x permutation test. **c,** Trial-by-trial peak rewarded DA transient amplitude (“rewDA”) from 1 session. Purple arrow and dashed line indicate the first rewarded (unstimulated) trial, whose amplitude is ∼0.065 dF/F—corresponding to the purple bar in the bar plot. Bar plot also shows mean peak rewDA amplitude for each stimulation condition and trial outcome (reward/no reward). Stimulated trials remained below the naturally occurring peak rewDA amplitude throughout the session and on average. **d-e,** Same analyses as in panels *a-b*, but for No-Opsin control mice. **f,** Stimulation fiber locations.

**Extended Data Fig 6.**
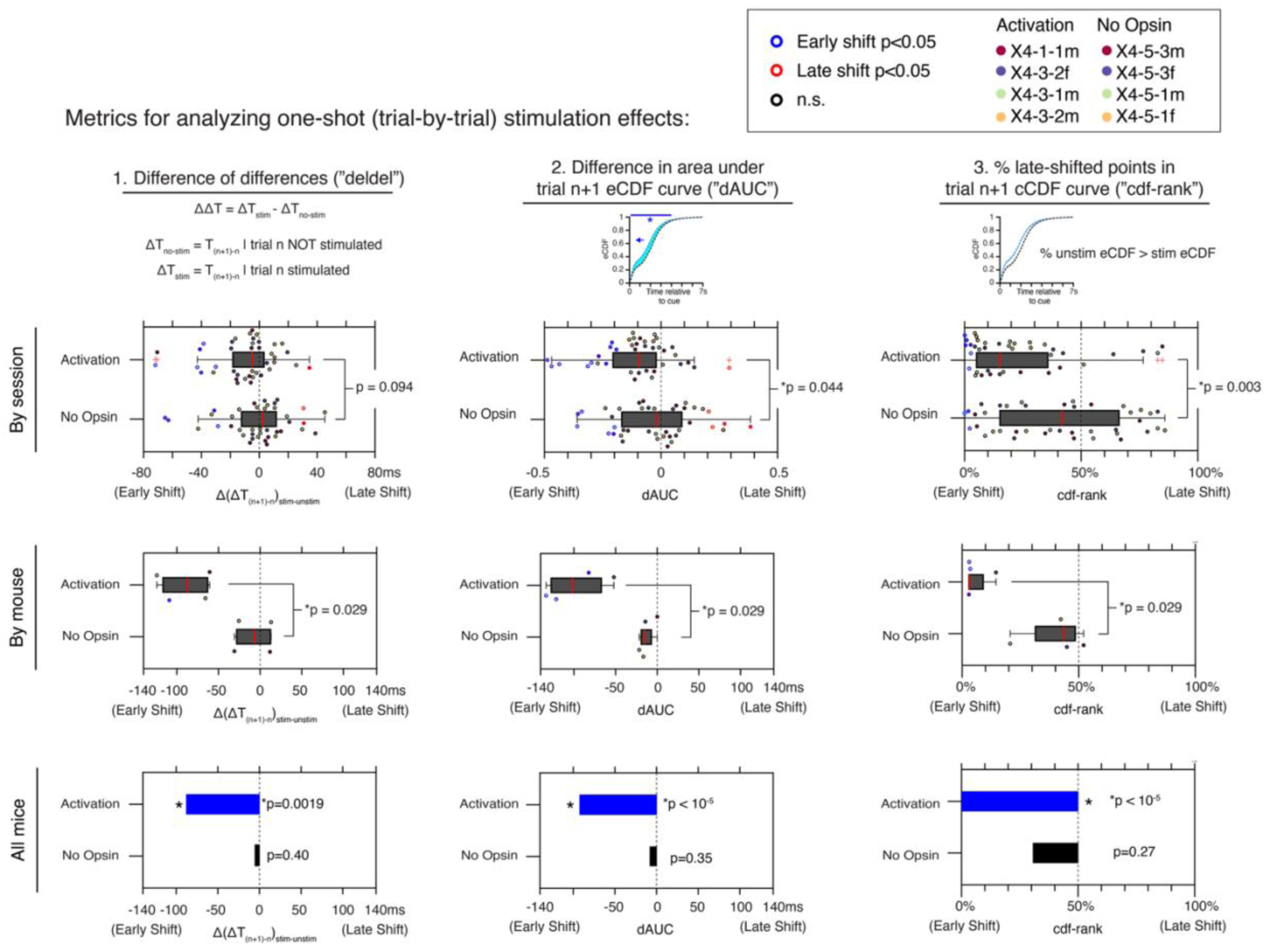
One-shot optogenetic manipulations—complementary analysis methods, all sessions and mice. Comparisons: stimulated vs unstimulated trials from the same (stimulated) session. Dot colors indicate whether the test statistic was significant at the individual session level (p<0.05). See Methods for detailed explanation of how each metric was derived.

**Extended Data Fig 7.**
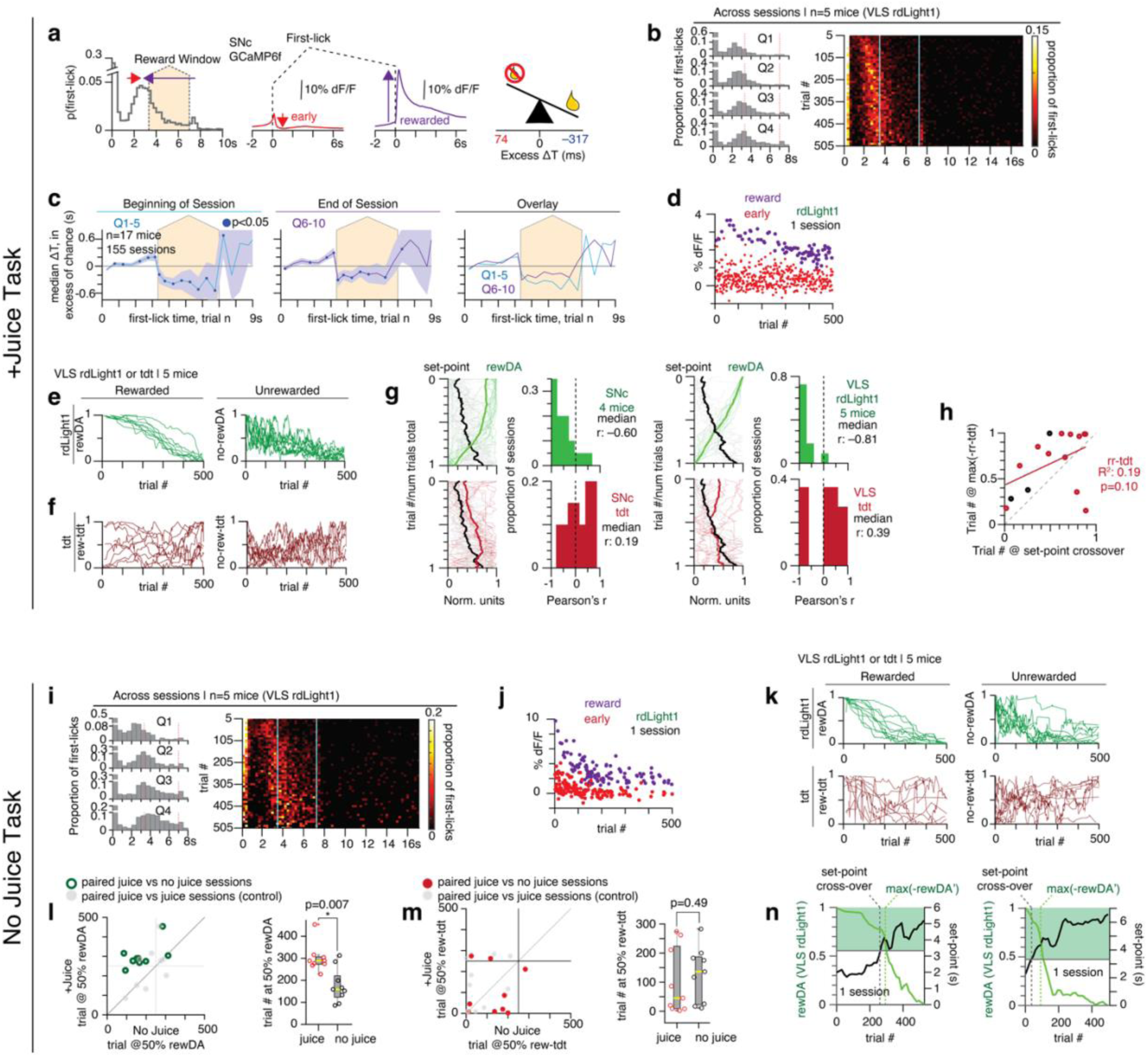
Timing behavior evolves over the course of each session. a, Skew in the timing distribution with respect to the reward window appears similar to asymmetry in outcome-related DA responses. **b,** Heatmap view of first-lick times over the course of the session, pooled across sessions and mice. The sessions shown were collected from the same mice as in panel *I* on paired sessions and are shown here for comparison. “Q1-4:” quartiles of trials in session, with Q1 representing the beginning of the session (i.e., trials 1-125). **c,** Timing updates—same analysis as Fig 1g, but calculated for trials at the beginning of the session (blue curves) vs. end of session (purple curves). To reduce non-stationarity of the timing distribution, sessions were divided into deciles of trials (Q1-10), and the timing update was then calculated as in Fig 1g. Fig 1g showed results from the whole session, where results from all ten deciles of trials were pooled. Here, pooled results from the beginning of the session (deciles 1-5) are shown separately from those from the end of the session (deciles 6-10). Right: overlay of beginning and end of session results. The effect of trial outcome on timing updates became less skewed toward the end of the session, attributable to rewarded trials being followed by smaller updates in Q6-10 (purple trace). **d,** Single-trial peak amplitude DA signal within 500ms after the first-lick shown over the course of one example session. By contrast to rewarded DA transients, unrewarded transients showed only a slight decrease in amplitude, and unrewarded timing updates (shown in panel *c*) likewise showed relatively little change across the session (beginning: +64ms, 95% CI: [+22ms, +103ms]; end: +94ms, 95% CI: [+35ms, +153ms]). **e,** Progression of single-trial DA transient amplitudes over the course of each session. Each line is one session. “rewDA:” reward-related DA peak amplitude; “no-rewDA:” unrewarded peak. **f,** Peak outcome-related tdTomato for the same sessions as panel *e*. **g,** Correlation of all single-session outcome-related DA (top) and tdt (bottom) transients with timing set-point. Thin traces: single sessions; thick traces: average. Left: SNc GCaMP6f; Right: VLS rdLight1 signals. Control rew-tdt signals are not consistently correlated with the timing set-point, arguing against a trivial correlation with ongoing body movements/optical artifacts. **h,** Comparison of the trial at which the tdTomato signal had its maximum negative derivative (max(-rew-tdt’)) vs. the trial at which the timing set-point (defined as the moving median of the first-lick distribution, as in Fig 4) first crossed into the reward window (“cross-over point”). Red points: +Juice sessions. Black points: No Juice sessions. The strong correlation of set-point cross-over with reward-related DA amplitude (**Fig 4e**) is not present in control tdTomato signals, arguing against a trivial correlation with ongoing body movements/optical artifacts. **i-k,** Same analyses as panels *b,d-f*, but from paired No Juice sessions from the same animals. **l-m,** Comparison of +Juice and No Juice Task. rewDA declines significantly earlier in the session in the No Juice Task compared to the +Juice Task, but there was no difference in the 50% max point for tdt data collected on the same sessions. Standard box plot with Wilcoxon rank sum test. The scatterplot points are the trial at which rewDA (panel *l*) or rew-tdt (panel *m*) reached 50% max peak amplitude for paired +Juice/No Juice session. Colored dots: the data for rdLight1 (green) or tdt (red); light grey dots: control data obtained from randomly pairing +Juice sessions from the same animal, which shows that the 50%-max transient amplitude tends to occur at about the same trial on sessions from the same task-variant when collected from the same mouse. **n,** Additional single-session examples of cross-over analysis in the No Juice task (analogous to **Fig 4e**).

**Extended Data Fig 8.**
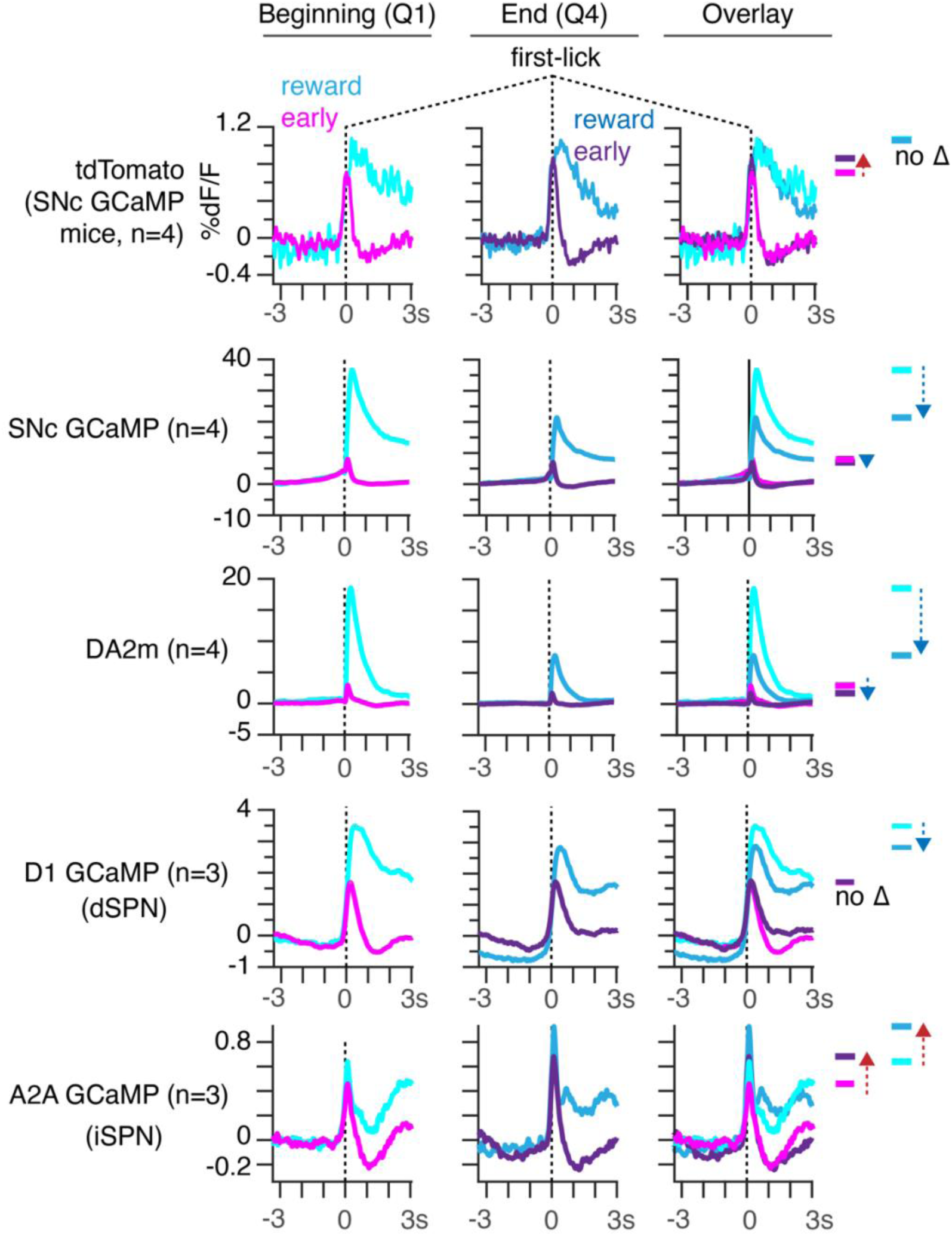
Comparison of outcome-related transient amplitudes between the beginning and end of the session. Q1: Beginning of session (first quartile of trials). Q4: End of session (fourth quartile). All DA/DAN recordings show decline in rewarded DA transient amplitude across the session, with comparably little change in unrewarded (early) transient amplitude. Optical artifacts (tdTomato) show the opposite direction of effect on unrewarded trials, whereas rewarded trial amplitude is similar across the session. As a comparison to the DAN and DA measurements, we also recorded GCaMP6f fiber photometry signals from striatal projection neurons (SPNs). Direct pathway SPNs (dSPNs) showed similar patterns to DA/DAN recordings. By contrast, indirect pathway SPNs (iSPNs) showed the *opposite* effect on average: both early-and reward-related iSPN transient amplitudes *increased* toward the end of the session. This difference is consistent with the putative sign of DA’s signaling interactions with SPNs as predicted from the DA receptors these cells primarily express: dSPNs primarily express excitatory metabotropic dopamine-1-receptors, whereas iSPNs primarily express inhibitory metabotropic dopamine-2-receptors. Moreover, the increase in iSPN GCaMP6f transient amplitude toward the end of recording sessions provides supporting evidence against bleaching-related artifacts as a possible explanation for the downtrend observed in reward-related DAN/DA signals (see **Methods**).

**Extended Data Fig 9.**
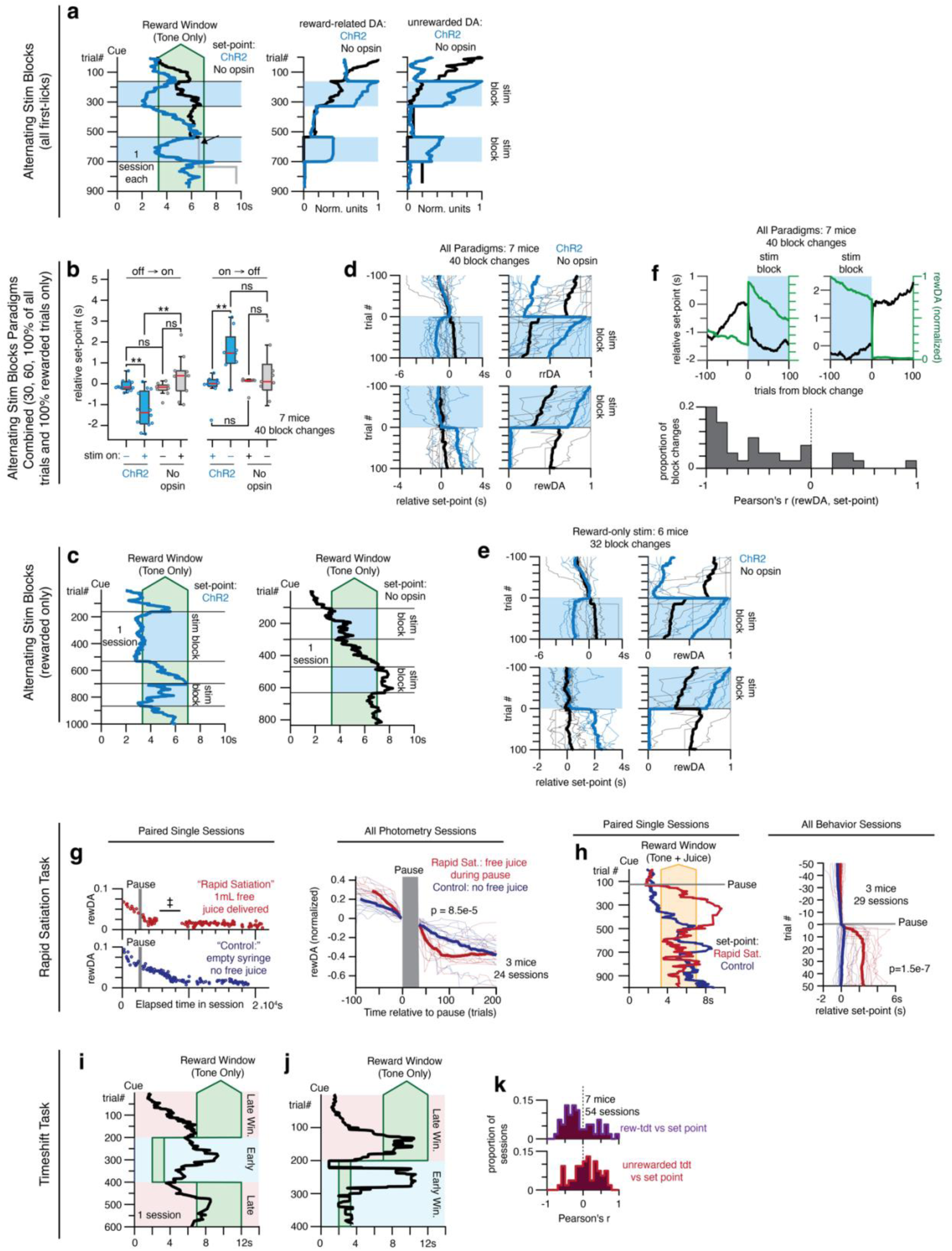
Timing set-point manipulations in more detail. a, Alternating Stimulation Blocks task (variant of task with all trials stimulated triggered on the first-lick during stimulation blocks). *Left*: Example single sessions (1 ChR2 mouse, 1 No Opsin mouse, 1 session each). Exposure to blue light initially caused a small early-shift in the No Opsin mouse, but this was much smaller than in the ChR2 mouse and did not recur during the second stimulation block. The No Opsin set-point trace is greyed after the arrow because the mouse did not participate on sufficient trials during blocks 4-5 to meaningfully calculate the moving median first-lick time. *Right*: Normalized outcome-related DA transient peak amplitude (“rewDA” or “unrewarded-DA”) in the example ChR2 session. Interestingly, a slow, gradual late-shift in the timing set-point was present even with ChR2 stimulation, and stimulated outcome-related DA-transient amplitudes declined progressively over the course of the session. **b,d,f,** Combined Alternating Stimulation Blocks results for all versions of this paradigm (Animals were stimulated triggered on either 30%, 60%, or 100% of first-licks during stimulation blocks (irrespective of trial outcome) or they were stimulated triggered only on first-licks that occurred during the reward window). ChR2: n=3 mice, 20 block changes; No Opsin: n=4 mice, 20 block changes. **b,** Same layout as **Fig 5d**, Wilcoxon rank sum, *p<0.05, **p<0.01, ***p<0.001. **c,** Comparison of ChR2 and No Opsin set-point dynamics, single sessions (2 mice, 1 session each). Despite exposure to blue LED, No Opsin mouse does not show early-shift during stimulation blocks. **d-e,** Single session block change set-point and rewDA dynamics. Thick trace: mean, thin traces: individual sessions. Blue: ChR2 mice, Black: No Opsin Mice. **f,** *Top:* Overlay of mean set-point and rewDA dynamics at block change. *Bottom:* Pearson’s correlation of set-point and rewDA across all Alternating Stimulation Block sessions. **g-h,** Rapid Satiation Task: comparison of set-point and rewDA dynamics for single sessions and across all sessions. **g**: Left: Single session examples shown for paired, interleaved sessions from the same mouse. Black line/dagger(‡) in the top single-session plot indicates that the interval without red dots had no rewarded trials during this particular session (the set-point was highly late-shifted with reduced participation during this time, see panel *h*). Right: All photometry sessions. Thin traces: single sessions; thick: average. **h**: Left: Behavior from the same paired single sessions as in panel *g*, Right: all behavior sessions. **i-j,** Timeshift Task: example No Juice session set-point dynamics. **k,** Unlike DA, the co-monitored tdt control channel was not consistently correlated with set-point changes and did not explain away the correlation between outcome-related DA transient amplitude and set-point (compare to **Fig 5l**).

**Extended Data Fig 10.**
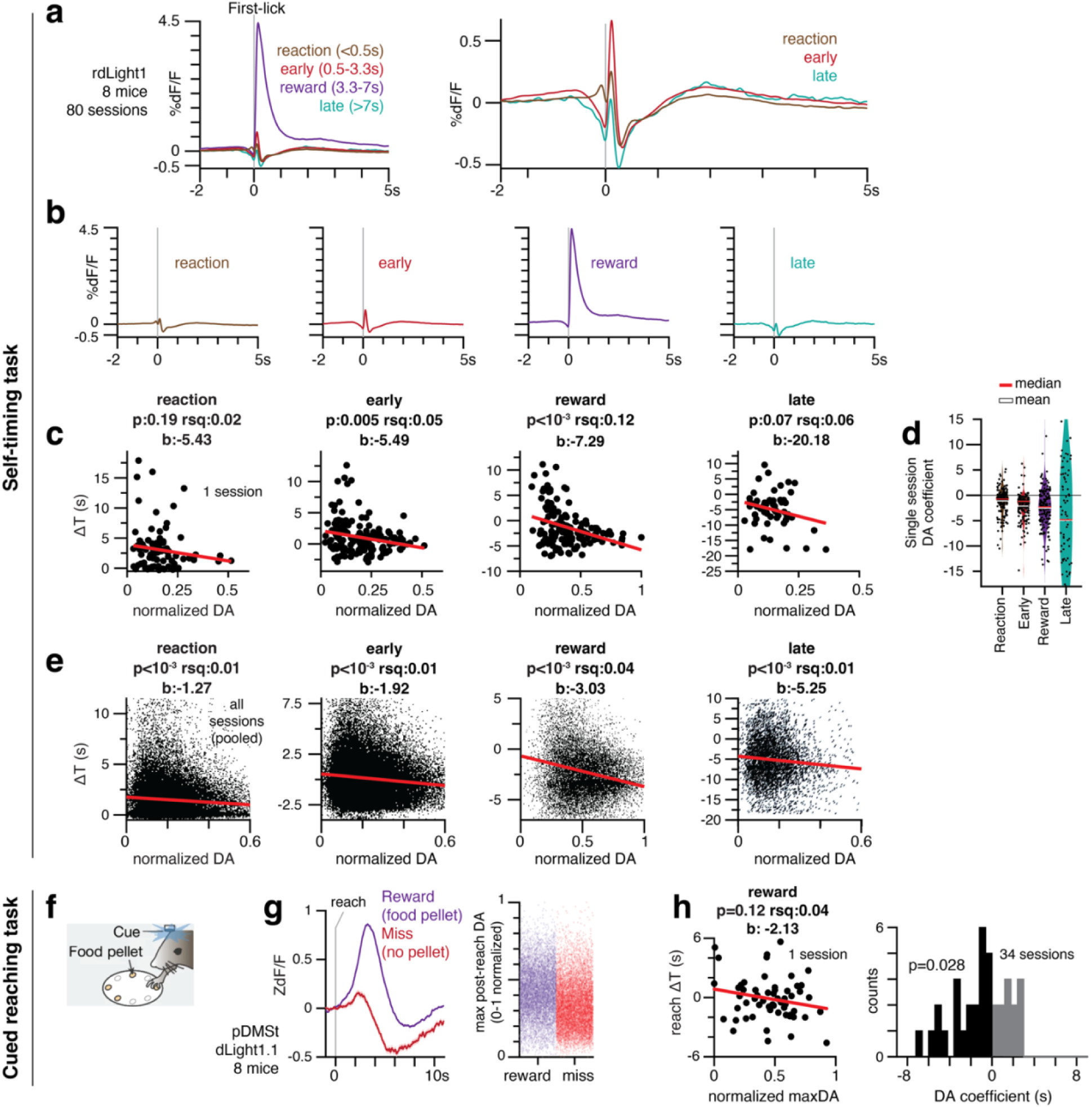
Outcome-related DA transient amplitude predicts a proportional change in movement time on the next trial. a-b, First-lick triggered average VLS DA signals for different trial outcomes, rdLight1. Rare late trial DA signals are similar to unrewarded early trials, but have lower amplitude transients (note that **Fig 1g** shows that these late trials are also associated with late-shifting in the median of the timing distribution on average (in excess of chance regression to the median effects), consistent with the observed DA dips and the GLM results). **c-e,** Simplified regression models for DA transient amplitudes, broken down by trial outcome (b: coefficient for DA predictor, which was encoded in the model as the mean amplitude of DA transient). DA transient amplitude predicts a proportional change in movement time on the next trial (inverse correlation) regardless of trial outcome. To avoid signal distortion, models were fit on actual change in movement time (ΔT), uncorrected for “regression to the median” effect, as in **Fig 2**. Note that **Fig 1g** (which addresses regression to the median) indicates that the negative ΔT observed after late/ITI trials was later-shifted than expected by chance. **c,** Single session example. **d,** Regression coefficients for all single sessions (all animals and signals (GCaMP and DA indicators). In most sessions, late trials were relatively rare and associated with high session-to-session variability in fit coefficient. To increase the number of late trials in the fit, we pooled sessions in panel *e* from the +Juice Task, late-window version of +Juice Task (see Methods), and Timeshift Task (179 sessions total). **e,** All sessions (all trials pooled). **f-h,** Cued reaching task, adapted from Reinhold et al., 2025.^63^ Mice received an optogenetic cue when a food pellet was available, and reaction times were measured as the latency between the cue and the reach. Interleaved catch trials omitted the food pellet (“miss”). Importantly, unlike the self-timed movement task, movement timing in the cued-reaching task had no bearing on whether reward was received. **g**, Reach-aligned striatal DA signals (dLight1.1) recorded from the posterior tail of the dorsomedial striatum (pDMSt). **h,** GLMs predicting trial-by-trial reach timing updates from DA and movement predictors were fit on rewarded trials within individual sessions. These models showed a similar trend to that in the self-timing task: higher peak DA amplitude predicted earlier reach times on the subsequent trial (p-value: 0.028, t-test). For miss trials, the distribution of DA coefficients was not significantly negatively skewed when the relationship was fit within single sessions (p=0.53), perhaps due to the relative rarity of miss trials. To increase statistical power, models were thus also fit on data pooled across sessions (1922 miss trials; 7786 reward trials), with rewarded and miss trials analyzed separately. In this pooled analysis, both trial categories showed significant effects consistent with those observed in the self-timing task (negative correlation between DA and timing updates; DA coefficient p-values for each model: Rewarded: p=0.030; Miss: p=0.038).

